# Influence of the extracellular domain size on the dynamic behavior of membrane proteins

**DOI:** 10.1101/2021.11.15.468619

**Authors:** Cenk Onur Gurdap, Linda Wedemann, Taras Sych, Erdinc Sezgin

**Author notes:** Contributed equally (Cenk Onur Gurdap, Linda Wedemann).

## Abstract

The dynamic behavior of plasma membrane proteins mediates various cellular processes such as cellular motility, communication, and signaling. It is widely accepted that the dynamics of the membrane proteins is determined either by the interactions of the transmembrane domain with the surrounding lipids or by the interactions of the intracellular domain with cytosolic components such as cortical actin. Although initiation of different cellular signaling events at the plasma membrane has been attributed to the extracellular domain (ECD) properties recently, the impact of ECDs on the dynamic behavior of membrane proteins is rather unexplored. Here, we investigate how the ECD properties influence protein dynamics in the lipid bilayer by reconstituting ECDs of different sizes or glycosylation in model membrane systems and analyzing ECD-driven protein sorting in lipid domains as well as protein mobility. Our data shows that increasing the ECD mass or glycosylation leads to a decrease in ordered domain partitioning and diffusivity. Our data reconciles different mechanisms proposed for the initiation of cellular signaling by linking the ECD size of membrane proteins with their localization and diffusion dynamics in the plasma membrane.

**SIGNIFICANCE STATEMENT:** We studied how the size and glycosylation of the proteins influences their dynamic behavior in a lipid bilayer by reconstituting the ECDs of different sizes or glycosylation in model membrane systems and analyzing their sorting into lipid domains as well as their mobility. We observe that increasing the ECD apparent mass leads to a decrease in membrane ordered domain partitioning and diffusivity. Our data reconciles multiple mechanisms proposed for the initiation of cellular signaling by linking the ECD properties of membrane proteins with their localization and diffusion dynamics in the plasma membrane.

## INTRODUCTION

The plasma membrane (PM) is a complex and dynamically heterogeneous system consisting of a lipid bilayer populated with various lipids and membrane proteins (MPs). About 40% of the human proteome is membrane-associated proteins (1) that are either integrally embedded into the PM or peripherally associated by the interactions with the headgroups of the inner leaflet lipids. Integral MPs are embedded in the membrane via one or multiple transmembrane domain(s) (TMD), with extramembraneous hydrophilic domains extending into the cytosol (intracellular domain, ICD) or into the extracellular environment (extracellular domain, ECD).

Cellular signaling events are often triggered at the plasma membrane via ligand-receptor interactions. For example, immune cell receptors are spatially reorganized and phosphorylated when interacting with their target ligands, eventually initiating a downstream signaling cascade. Multiple mechanisms are proposed to explain the initial protein reorganization that leads to the activation (phosphorylation) of the receptors. Partitioning of signaling proteins into membrane domains (2), their ECD size (3–5), their allosteric interactions with lipids (6), and their mobility altered by the ligands (7, 8) are some examples of these mechanisms. Although there is compelling evidence for each of these mechanisms for immune signaling, the dominating mechanism is still unclear. In this work, we investigated whether these seemingly unrelated mechanisms can be in fact intertwined. To this end, by using model membranes and fluorescence correlation spectroscopy (FCS), we investigated the relationship between the ECD size or glycosylation of various immune signaling proteins, their molecular diffusion, and their domain partitioning.

Model membranes are widely used as a synthetic proxy to investigate biophysical aspects of the PM, such as membrane domain formation (9). In these simple model membrane systems, saturated lipids form a liquid-ordered phase (L_o_) with the help of cholesterol, and unsaturated lipids form a liquid-disordered phase (L_d_). Different artificial or cell-derived model systems are available to study the phase separation phenomenon (9). Giant unilamellar vesicles (GUVs) and giant plasma membrane vesicles (GPMVs) are commonly used cell-sized spherical free-standing lipid bilayers. While GPMVs are cell-derived and thereby inherently more complex in lipid and protein compositions, GUVs are artificial systems with finely controlled composition. Both systems could display the co-existing phases to mimic the lateral heterogeneity of the PM (10).

Proteins were shown to partition into one of these lipid phases preferentially. Such selective protein partitioning can induce lipid environment-dependent conformational change and promote lipid-driven protein associations, crucial for protein function, such as immune cell function and viral dynamics (11–14). Recent studies showed that TMD length, lipid accessible surface area of TMD, and the addition of fatty acid molecules such as palmitoylation at the TMD can influence protein partitioning in membranes (15–17). Glycosylphosphatidylinositol (GPI)-anchored proteins partition based on the structure of their lipid anchors (18). Critical factors in phase separation and protein partitioning are reviewed in detail in refs (19, 20). While there is extensive research on the influence of TMD properties on phase partitioning of proteins (17), the role of ECD properties (e.g., length, molecular weight, and glycosylation) on partitioning is largely unexplored. Some of the earlier work performed using biochemical methods that detergent resistance suggested a non-existing role of ECDs in membrane domain partitioning of proteins, such as CD4 (21). In recent years, however, the role of ECDs on the membrane structure and dynamics have been addressed using state-of-the-art methods. For example, it was shown that protein assemblies at the extracellular side of the membrane induce membrane reorganization, such as membrane bending (22), tube formation (23), concomitant polymerization (24), or domain dissolution (25). Moreover, ECD size has been one of the major players in immune synapse formation (3), continuously being studied in the context of lipid remodeling (26). Finally, different lengths of DNA oligos showed different phase partitioning in phase-separated synthetic vesicles (27). Altogether, this evidence necessitates a thorough study on the effect of ECDs on the partitioning behavior of the MPs.

To exhibit their function, MPs need to be dynamic since their lateral mobility allows them to interact with other proteins and thereby form complexes. Therefore, measuring the lateral mobility of proteins is crucial to gain a mechanistic understanding of their function. Influenced by the compositional complexity, the dynamics of MPs are not only governed by simple Brownian motion, but it is rather more complex, mediated by lipid domains, protein-protein interactions, and cytoskeletal elements (28–34). According to the established Saffman and Delbrück model, when a molecule is anchored to the membrane, its diffusion depends on membrane viscosity (η_m_), membrane thickness (h), surrounding bulk fluid viscosity (η_f_) and weakly on the radius of the TMD (R) (k_B_ is the Boltzman constant, γ is the Euler’s constant and T is temperature) (35):

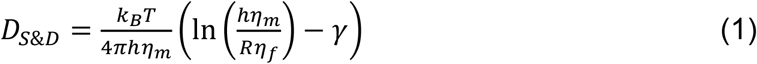

There have been several corrections to this model (36–43). However, these corrections are mainly focused on the impact of lipid environment, TMD, and ICD of the proteins on lateral diffusion, with the exception of GPI-anchored proteins, where some hints of the influence of extramembraneous size on diffusion were observed (44, 45). Recently, *Houser et. al*. suggested a correlation between the ECD molecular weight and diffusion under crowded conditions, showing that larger proteins display a slower diffusion due to larger area coverage (crowding) (46). Furthermore, computational simulations (47) and experimental methods (48) have predicted that the diffusion of MPs might also be influenced by their ECDs. Therefore, it is crucial to reveal how ECD properties influence the diffusion of MPs and to develop new models taking this role into consideration.

In this work, we reconstituted polyhistidine-tagged ECDs of different sizes or glycosylation in model membranes functionalized with nickel-chelating lipids. In this setting, TMDs and ICDs are absent, hence the impact of ECD properties on protein behavior can directly be elucidated. To monitor the phase partitioning of ECDs, we used phase-separated GUVs, and interestingly, we found that protein sorting to the L_o_ phase decreases with increasing ECD mass. Furthermore, we studied ECD mobility in GUVs of different compositions and GPMVs. We found that protein mobility also decreases as the mass of the ECD increases for smaller molecules but not for large proteins. These changes correlate perfectly with the apparent mass of the proteins rather than their predicted mass by amino acid sequence. Finally, our data showed that glycosylation is a critical factor that influences both partitioning and diffusion of proteins. When glycosylation was removed from proteins, they exhibited higher Lo partitioning and faster diffusion. Overall, our data suggest a critical role of the ECD properties on membrane protein behavior in the plasma membrane and paves the way to a complete understanding of membrane protein dynamics that also takes interaction with the extracellular matrix and glycocalyx into account.

## MATERIALS AND METHODS

### Lipids, proteins, fluorescent probes

1-Palmitoyl-oleoyl-sn-glycero-phosphocholine (POPC), 1,2-Dipalmitoyl-sn-glycero-3 phosphocholine (DPPC), 1,2-Dioleoyl-sn-glycero-3-phosphocholine (DOPC), sphingomyelin (SM), cholesterol (Chol), 1,2-Dioleoyl-sn-glycero-3-[(N-(5-amino-1-carboxypentyl)iminodiacetic acid)succinyl] (nickel salt) (18:1 DGS-NTA(Ni)), and 1,2-Dipalmitoyl-sn-glycero-3-[(N-(5-amino-1-carboxypentyl)iminodiacetic acid)succinyl] (nickel salt) were obtained from Avanti Polar Lipids. Lipid stocks were stored in chloroform under nitrogen at -20 °C.

ECD fragments (modified C-terminally with 6xHis tag) of cluster of differentiation (CD) 59, (Cat.# 12474-H08H), CD2 (Cat.# 10982-H08H), intercellular adhesion molecule (ICAM-1) (Cat.# 10346-H080H), CD45 (Cat.# 16884-H08H), and CD34 (Cat.# 10103-H08H) were purchased from SinoBiological. ECD fragments (modified C-terminally with 10xHis tag) Podocalyxin (PODXL) (Cat.# 1658-PD) and PODXL2 (Cat.# 1524-EG) were obtained from R&D Systems. The ECD fragments were dissolved in ultrapure water to a concentration of 0.5 mg/mL and aliquoted in 50 μL, snap-frozen, and stored at -80°C. For immediate usage, protein solutions were stored at 4°C avoiding freeze/thaw cycles. 6xHis-tagged Alexa 488 (A488-6xHis), 1.2 μg/mL, was custom-made by Cambridge Research Biochemicals. Abberior STAR Red-1,2-Dipalmitoyl-sn-glycero-3-phosphatidylethanolamine (AbStR-DPPE) obtained from Abberior GmbH.

For protein labeling, Alexa Fluor 488 NHS Ester (A488) was purchased from Thermo Fisher and performed according to NHS labeling protocol. Briefly, dye solution in DMSO (concentration not lower than 10 mg/ml) was added to protein solution in pure water (0.5 mg/ml; 1 mg/ml for CD34, PODXL, and PODXL2) and incubated for 1 hour at 300 rpm in the presence of 100 mM NaHCO_3_ (pH 8.3). Molar excess of dyes for each protein is displayed in Supp. Table 1. Purification was performed using spin columns (7 kDa exclusion limit, 0.5 mL Zeba spin desalting columns, ThermoFisher Scientific). After purification, the protein concentration (C_p_ final) was measured via nanodrop with the knowledge of the individual extinction coefficients, estimated from the Uniprot amino-acid sequences using the ProtParam tool by Expasy **(Supp. Table 1)**.

For deglycosylation treatment of the proteins, His-tagged CD34, PODXL, and PODXL2 were mixed with Protein Deglycosylation Mix II (New England Biolabs, #P6044S). The suggested protocol by the supplier was used with the exception of incubation time (the proteins were only incubated at RT for 20 h).

### Preparation of model membranes

GUVs were produced via the electroformation method. A lipid mixture was dissolved in chloroform-methanol (1 mg/mL POPC, POPC:Chol (1:1), DPPC:Chol (1:1), or DOPC:SM:Chol (2:2:1, phase-separated) with further addition of 1 mol % of Ni-NTA was spread on platinum wires and left for the solvent to evaporate. It was then dipped into 300 mM sucrose solution in a custom-build, Teflon-coated electroformation chamber. Electroformation was performed for 1 h with 10 Hz AC field (2V) followed with 30 mins in 2 Hz (DG822, Rigol) for vesicle release from the wires. For the DPPC and DOPC:SM:Chol mixtures, an external heat source was used at 55 °C for the electroformation, and the produced GUVs were then slowly cooled down to RT.

GPMVs were isolated from CHO (Chinese Hamster Ovary) cells which were grown to about 70% confluency in Dulbecco’s Modified Eagle Medium media supplemented with 10% fetal bovine serum (FBS, Sigma) and 1% L-glutamine, or from HeLa cells with about 70% confluency (in DMEM supplemented with 10% FBS and 1% L-glutamine). Cells were prepared two days before the experiments by seeding them onto 6-well cell culture plates. They were then washed twice with 2 mL GPMV buffer (10 mM HEPES, 2 mM CaCl2, and 150 mM NaCl, pH 7.4) and incubated in 1mL GPMV buffer for 2 h. To form vesicles from CHO cells, the buffer was additionally supplemented with 25 mM paraformaldehyde (PFA, Sigma) and 2 mM 1,4-Dithiothreitol (DTT, Sigma). To form phase-separated vesicles, HeLa cells were used to isolate GPMVs, where GPMV buffer was supplemented with 25 mM PFA and 25 mM DTT.

To insert Ni-NTA lipids into GPMVs, the supernatant containing GPMVs were then extracted and transferred into a new Eppendorf tube. In another Eppendorf tube, the chloroform from 20 μg of Ni-NTA lipid (5 mg/mL stock concentration) was evaporated under a stream of nitrogen. First, 1 μL of PBS and then 180 μL of extracted GPMVs were added onto the dried Ni-NTA lipid. After vortexing for 10 seconds, 65° heat was applied for 5 mins, followed by cooling for 10 mins to the RT. The mixture was then left to sit at RT for 20 mins before labeling. In phase-separated GPMVs, due to the high concentration of DTT treatment to induce phase separation, the nickel on the lipid was likely reduced and sometimes formed a brownish pellet at the bottom of the tube. However, the formation of pellets from the Ni-NTA lipid was not the issue for GPMVs prepared from CHO cells, mainly because of 10 times less concentrated DTT treatment during its extraction.

For labeling both GUVs and GPMVs, AbStR-DPPE at a concentration of 0.25 μg/mL was added into the vesicle solution and incubated for 15-30 mins to allow its integration into the membrane. Samples were further supplemented with different A488-labeled ECDs with a final concentration of 0.5 μg/mL and incubated for 15-30 mins for His-Ni-NTA coupling. To reduce the background signal by unbound proteins, the labeled GUVs can be subsequently washed twice with 750 μL PBS, followed by allowing sedimentation for 20-30 mins after each wash. The 80-100 μL of GUVs from the bottom of the tube after sedimentation can then be used for measurements. The confocal imaging was performed in Ibidi 8-well plastic bottom chambers, while Ibidi 8 or 18-well glass-bottom chambers (#1.5H) were used for the FCS measurement. The wells were pre-treated with 1 mg/mL bovine serum albumin beforehand for at least 1 h and washed three times with PBS. All pipetting was performed gently, and pipette tips were cut to reduce shear stress on the vesicles.

### Confocal imaging of phase-separated GUVs

The Zeiss LSM 780/980 confocal laser scanning microscope equipped with a 40x 1.2 water-immersion objective was used for imaging and FCS experiments. The microscope’s objective was additionally provided with a DIC Prism (DIC Prism III PA 63x/1.40, model 426957, Zeiss) for excitation laser depolarization for imaging experiments. Green and far-red fluorescence were excited with 488 nm and 633 nm lasers, and their fluorescent emission was detected in two channels of the 32-channel detector within the spectral windows of 498–580nm and 641–696 nm, respectively. The pinhole parameter was set to 1 Airy Unit. The dynamic range was 12 bits (4096 grey value units), with four times line averaging. The confocal image was taken by positioning the focal plane at the GUV equatorial plane. Line plot profile of fluorescence intensity was performed using ImageJ (49). Fluorescence intensities in L_d_ and L_o_ phases were calculated from the plot peaks. Two directly opposite sides were chosen in the vesicles to eliminate the light intensity difference due to laser polarization. L_o_ partitioning (%L_o_) was calculated by dividing the fluorescence intensity of L_o_ by the sum of the fluorescence intensities of L_o_ and L_d_.

### FCS measurement

The calibration of the pinhole position and the correction collar of the objective for FCS was performed before the measurement using Alexa 488-Alexa 647 solution mixture in water in the same Ibidi chamber as the samples. For FCS in GUVs or GPMVs, the focal plane was positioned at the bottom or the top of the vesicle, respectively. FCS curves were obtained for 5-10 seconds with low laser power (2-5 microwatts) with 5-10 repetitions to prevent photobleaching. The obtained correlation curves were fit and analyzed by the FoCuS-point software (50).

### ECD conformation prediction

The three-dimensional conformation of the ECDs were in situ predicted via AlphaFold (AlphaFold2_advanced.ipynb, collabfold). The details of the parameter are shown in Supp. Table 2, and the amino acid sequences are shown in Supp. Table 3. These structures were subsequently aligned in UCSF ChimeraX (Version 1.3) to published AlphaFold predictions of the full-length proteins to assess the folding of the isolated ECDs.

### Statistical analysis

The statistical analysis of each data set is explained in corresponding figure legends.

## RESULTS AND DISCUSSION

We used GUVs and GPMVs as model systems to mimic cellular membrane surfaces and study protein behavior. The main advantage of GUVs is finely controlled lipid composition while that of GPMVs is the near-native composition. We studied protein phase partitioning in phase-separated GUVs, and protein diffusion in GUVs and GPMVs.

### Saturated and unsaturated nickel-chelating lipids partition into Lo and Ld phase, respectively

An important dynamic behavior of proteins is their partitioning to certain lipid environments. To study the role of ECD size on protein partitioning, we first prepared phase-separated GUVs (DOPC/SM/Chol, 2:2:1) with 1 mol% of nickel-chelating lipid that can directly bind to 6xHis-tagged ECDs **(Fig. 1A)**. To visualize phase separation, we used AbStR-DPPE that partitions preferentially to L_d_ **(Fig. 1A, B)**. To evaluate the partitioning of proteins in both ordered and disordered phases, we used two different nickel-chelating lipids: unsaturated 18:1/18:1 (we will refer to it as di-oleyl (DO) Ni-NTA) and saturated 16:0/16:0 Ni-NTA (we will refer to it as di-palmitoyl (DP) Ni-NTA). To recognize and characterize different Ni-NTA lipids, we used Alexa 488 labeled 6xHis (A488-6xHis). In GUVs with DO Ni-NTA, A488-6xHis preferentially binds to L_d_, and in GUVs with DP Ni-NTA, it is preferentially associated with L_o_ **(Fig.1B)**.

**Fig. 1.**
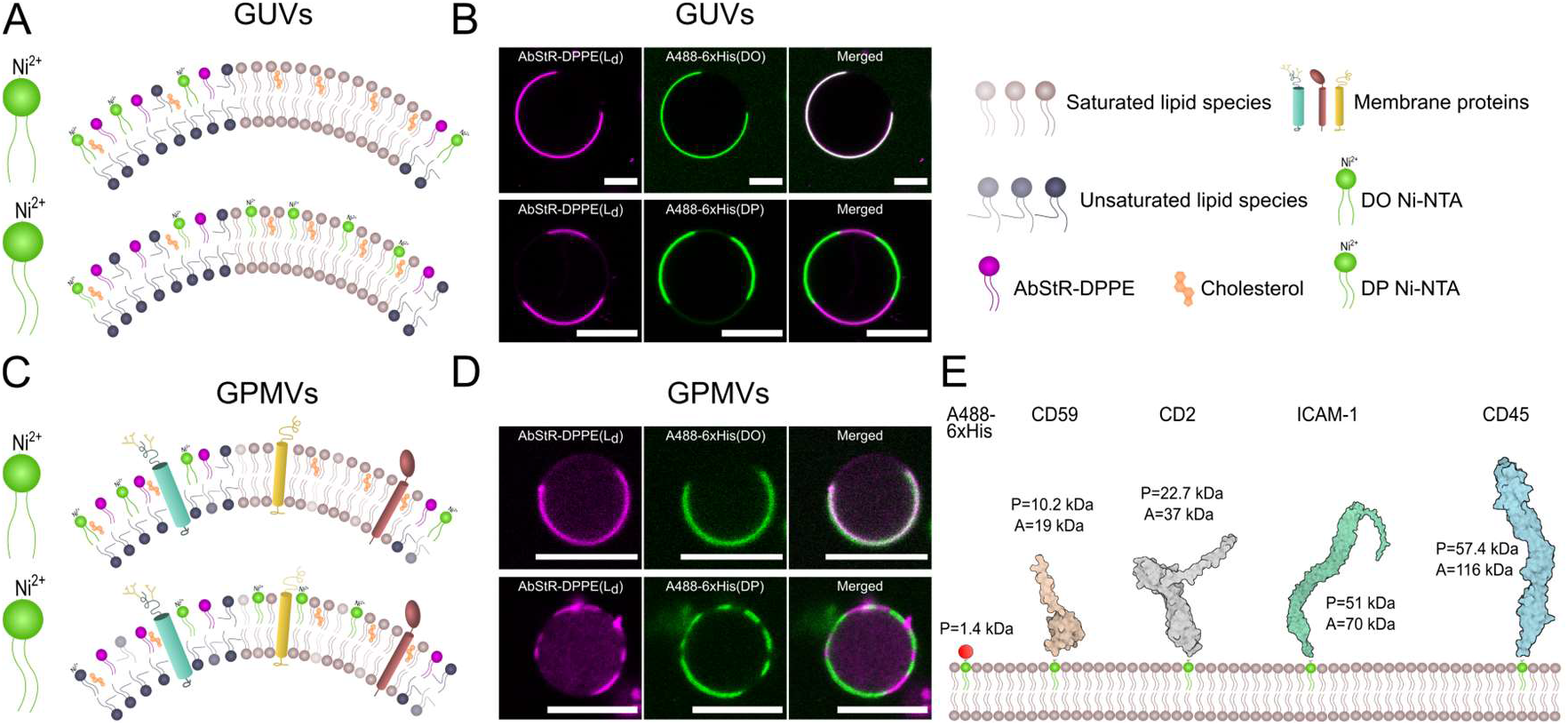
Ni-NTA lipid constituted biomimetic membranes with His-tagged proteins. (A) Cartoon and (B) confocal microscopy images of phase-separated GUVs incorporated with DO and DP Ni-NTA. GUVs were composed of the ternary mixture of DOPC/SM/Chol (2:2:1). (C) Cartoon and (D) confocal microscopy images of phase-separated GPMVs incorporated with DO and DP Ni-NTA. GPMVs were extracted from HeLa cells. AbStR-DPPE (magenta) and DO Ni-NTA incorporate preferentially in the disordered phase, whereas DP Ni-NTA prefers the ordered phase. Scale bars are 10 µm. (E) Proteins of interest used in the study (sizes are representative based on the structures). It shows the predicted (P) and apparent (A) mass of the proteins based on amino acids sequence and a SDS page, respectively.

Artificial phase domains in GUVs display high contrast in membrane order between L_o_ and L_d_ domains. Alternatively, cell-derived GPMVs present more complex domains with a less dramatic difference in membrane order. To evaluate the partitioning of these Ni-NTA lipids in a more natural system, we incorporated nickel-chelating lipids into GPMVs **(Fig. 1C)**. However, incorporating Ni-NTA lipids to GPMVs is rather challenging compared to GUVs presumably due to the charged surface of GPMVs (as the negatively charged lipids flip from the inner leaflet). Another possible reason might be the high DTT concentration used to form phase-separated GPMVs, which possibly reacts with nickel and reduces it. Despite these challenges, we obtained a small number of phase-separated GPMVs with nickel-chelating lipids attached to A488-6xHis and confirmed that DO Ni-NTA partitions to the disordered domains whereas DP Ni-NTA prefers the ordered domains **(Fig. 1D)**.

Since GUVs with Ni-NTA lipids were significantly easier to prepare, we continued the rest of the partitioning experiments solely using GUVs, where we evaluated the role of ECD size on partitioning. For this purpose, we selected ECDs of proteins that are mainly involved in immune signaling: CD59, CD2, ICAM-1 and CD45, whereas A488-6xHis was used as a control **(Fig. 1E)**. Predicted (P) and apparent (A) molecular mass of these proteins are shown in Fig.1E, and the number of amino acids in these peptides are shown in Supp. Table 1. P and A are different primarily due to the glycosylation of the proteins. P is the molecular weight calculated using the amino acid sequence solely while A is measured in a SDS page under reducing conditions, taking glycosylation of the protein into account. A is larger than P for all the proteins we used **(Fig. 1E)** since they are produced in HEK293 cells and preserve native mammalian glycosylation patterns. The difference between A and P shows the level of glycosylation; the larger the difference, the more glycosylation the protein has.

### ECD size influences domain partitioning

ECDs do not often directly interact with the PM. However, given the recent evidence on the impact of extracellular protein organization on membrane domains (22–25), ECDs might play a role in protein compartmentalization in the membrane. Earlier work using detergent resistance suggested certain sequences in ECDs can be ordered domain targeting sequences (51–53). However, the size of the ECDs has not been evaluated in this context. Therefore, we set out to evaluate whether ECD size affects protein partitioning in phase-separated GUVs. To investigate this, ECDs of different sizes **(Fig. 1E)** were incorporated into phase-separated GUVs **(Fig. 2)**. The fluorescence intensity of the two peaks in the line profile across the GUV image (A488-protein and AbStR-DPPE) was used to calculate the percentage of L_o_ partitioning **(Fig. 2A)**.

**Fig. 2.**
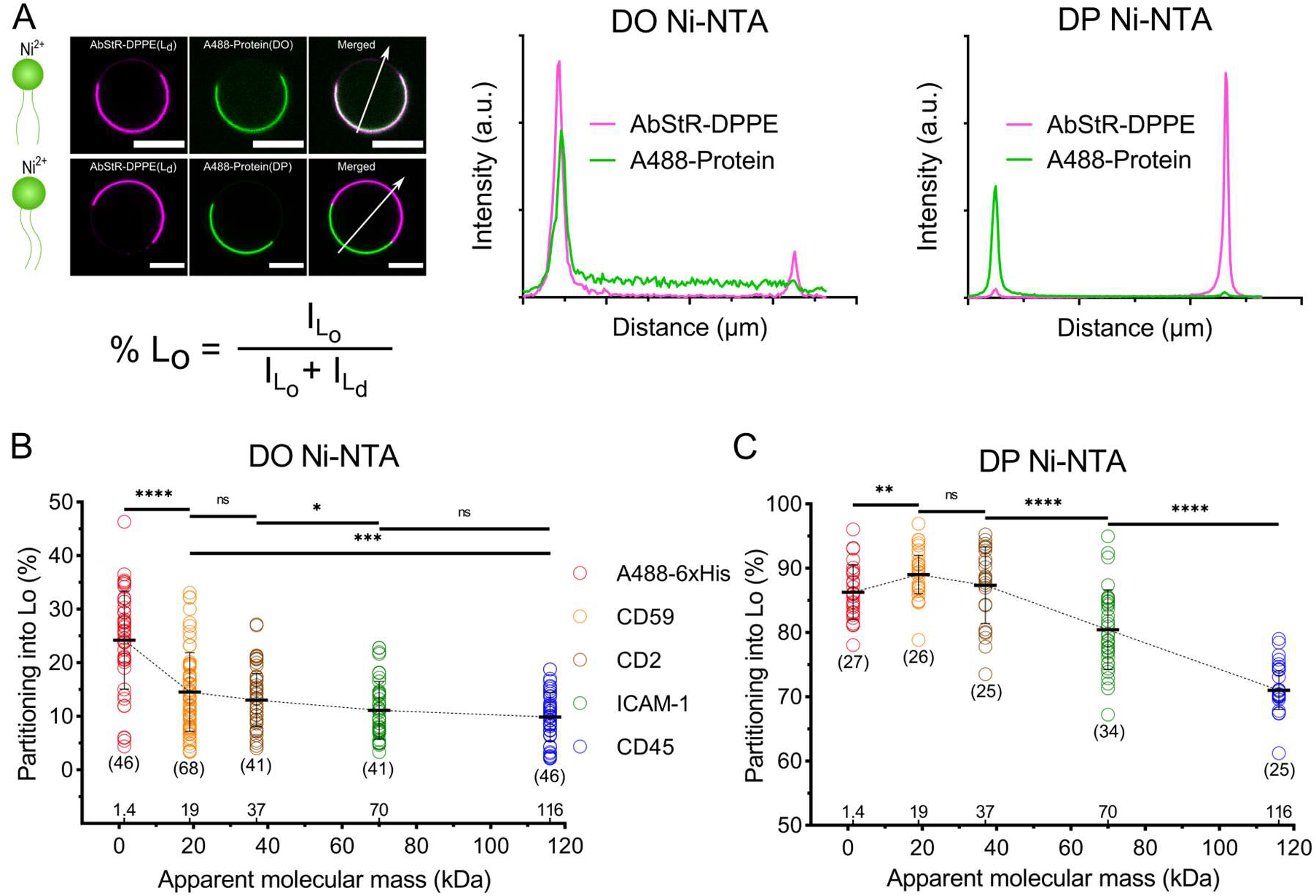
ECD mass plays a role in the protein partitioning in GUVs. (A) Line profile of fluorescence intensity across the GUVs indicated by the white arrow in the confocal images was used to calculate L_o_ percentage with the equation shown below the images. Scale bars are 10 µm. (B) L_o_ partitioning of different ECDs attached to DO Ni-NTA in GUVs. Student’s t-test (two-tailed, non-parameterized) was used to determine significance (****P<0.0001; ***P=0.0008; *P=0.0393; ns: non-significant). (C) L_o_ partitioning of the different ECDs attached to the DP Ni-NTA lipid in GUVs. Student’s t-test (two-tailed, non-parameterized) was used to determine significance (**P=0.005). Horizontal lines show the mean and error bars represent the standard deviation. The numbers of data points obtained from experiments are indicated on the graphs in parentheses.

To examine whether the concentration of the proteins on the surface affects partitioning, we first performed a control experiment with varying protein concentrations within the ranges we used (0.5 µg/ml up to 2 µg/ml). The result indicates that there is no effect of the concentration of the largest protein on its partitioning preference within the investigated concentration range **(Supp. Fig. 1)**. It is worth noting that, at higher concentrations, protein crowding is likely to play a role; however, we always used a low concentration of proteins for our study, avoiding the impact of crowding effects on our investigation.

We next investigated the effect of ECD size on protein partitioning. As mentioned before, ECD size can be estimated using the amino acid sequence (predicted molecular mass), which ignores the post-translational modifications such as glycosylation. Since many membrane proteins are highly glycosylated, the apparent molecular mass is generally higher when measured under reduced conditions. Therefore, we first measured the partitioning as a function of apparent molecular mass; however, the predicted mass and apparent mass for these proteins show the same increasing trend, i.e., as the predicted mass increases, the apparent mass also increases for the chosen proteins. As expected, in GUVs with DO Ni-NTA, proteins partitioned preferentially to L_d_, whereas in GUVs with DP Ni-NTA to L_o_. Interestingly, in both systems, L_o_ partitioning decreased as the apparent molecular mass increased **(Fig. 2B, C)**. These findings imply that apparent ECD mass is one of the determining factors for protein partitioning, and larger (and potentially highly glycosylated) proteins might be less abundant in ordered domains. This could be explained by different scenarios that need to be addressed. First, we needed to rule out any direct and differential interactions between different ECDs and the membrane. To this end, we generated the structures of the proteins via AplhaFold **(Supp. Fig. 2, Supp. Table 2, 3)** and measured the propensity of different proteins to interact with the membrane **(Supp. Fig. 3)** (54). This analysis showed that the hydrophobicity of the proteins is very similar, and there are not any notable potential interactions between the membrane and the ECDs. Therefore, the differences in hydrophobicity or membrane interactions cannot be responsible for the size-dependent partitioning we observed. Clustering might be another possible reason for different partitioning. To test whether our proteins clustered differently, we measured the apparent brightness of each protein with FCS. Apparent brightness for proteins was random (i.e., did not follow any trend with the size of the protein) and likely determined by the number of conjugatable amino acid moieties in the ECDs **(Supp. Fig. 4)**. To be able to account for the reason behind the discrepancy in the partitioning of ECDs of different sizes, we needed to add more proteins into our measurements. While the proteins we tested above have a clear trend in apparent mass correlated with their partitioning, they also have a similar trend in predicted mass. To pinpoint the mechanism causing the difference in partitioning, we tested three more proteins with different trends in predicted vs. apparent mass.

### ECD glycosylation influences domain partitioning

Glycosylation is a posttranslational modification, leading to an addition of carbohydrates on proteins. It makes the proteins more rigid and heavier such that highly glycosylated proteins can be a few times heavier than their predicted mass (from their amino acid sequence). To test how the discrepancy between predicted and apparent mass affects the partitioning of ECDs, we picked three proteins from the same family with different degrees of glycosylation; CD34, PODXL, and PODXL2 **(Fig. 3A)**. PODXL is the most heavily glycosylated form, followed by PODXL2 and CD34. Interestingly, the predicted mass of PODXL is lower than PODXL2, but due to glycosylation, the apparent mass of PODXL becomes significantly higher than PODXL2. Therefore, the partitioning behavior of these three proteins will show whether apparent or predicted mass determines the partitioning. When we compared their partitioning in GUVs, we observed a clear trend between partitioning and the apparent mass but not the predicted mass **(Fig. 3B, C)**. To further confirm the role of glycosylation in partitioning, we removed the sugar residues from the proteins by a deglycosylation enzyme mixture (55). After the enzyme treatment, we observed an increase in the partitioning of proteins into Lo, except CD34, confirming the role of glycosylation in partitioning **(Fig. 3D)**. CD34 is the least glycosylated of these three proteins, therefore it is expected to see the smallest difference for CD34 upon deglycosylation. However, seeing no difference is presumably due to incomplete effect of the enzyme on different glycosylation patterns (N-, O-glycosylation or sialylation).

**Fig. 3.**
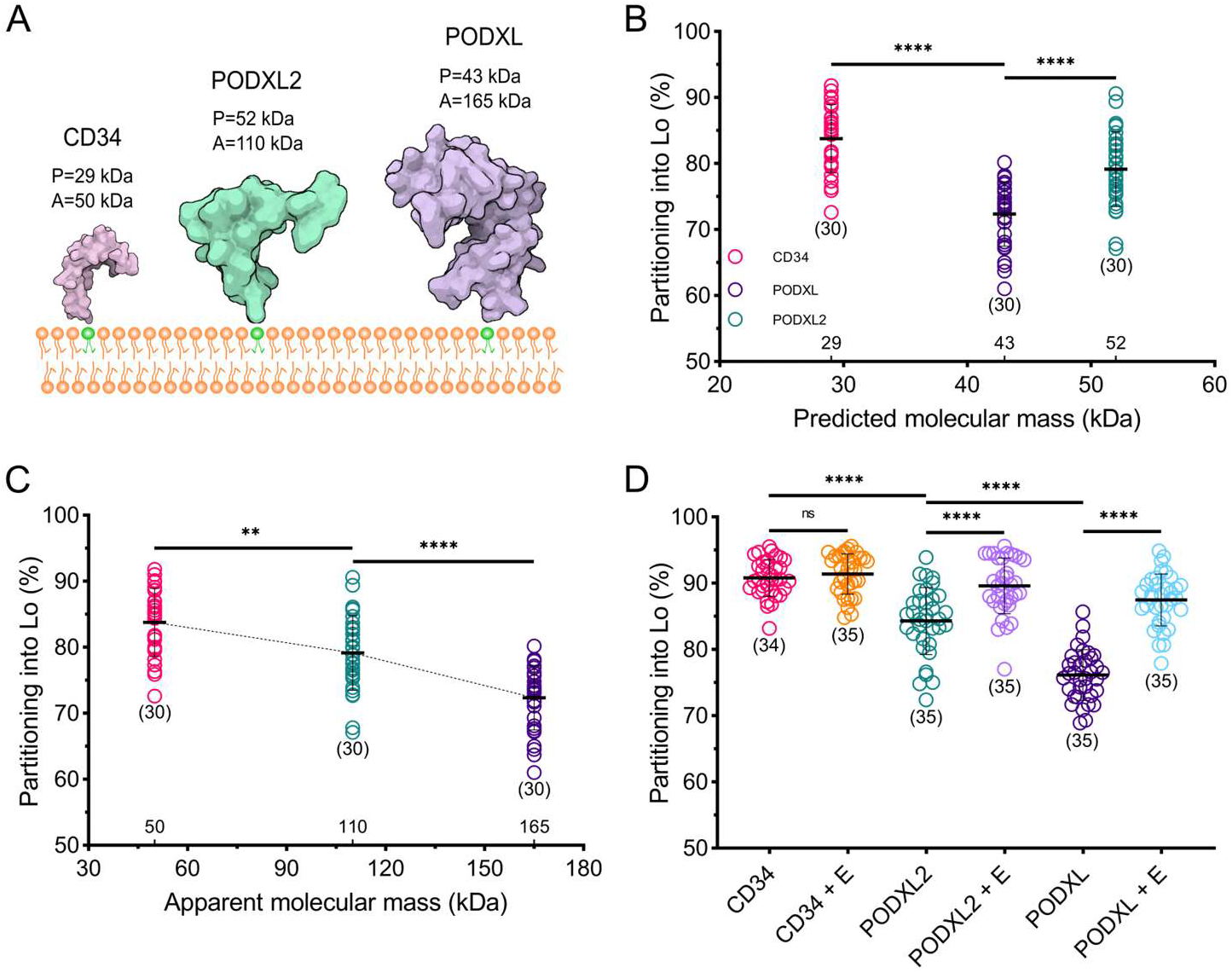
Glycosylation and apparent ECD mass play a role in protein partitioning in GUVs. (A) Cartoon of CD34, PODXL and PODXL2 (sizes are representative based on the structures). (B) L_o_ partitioning of different ECDs attached to DP Ni-NTA in GUVs as a function of the predicted mass. Student’s t-test (two-tailed, non-parameterized) was used to determine significance (****P<0.0001) (C) L_o_ partitioning of the different ECDs attached to the DP Ni-NTA lipid in GUVs as a function of the apparent mass. Student’s t-test (two-tailed, non-parameterized) was used to determine significance (**P=0.0028). (D) L_o_ partitioning of the different ECDs after the deglycosylation enzyme treatment. Student’s t-test (two-tailed, non-parameterized) was used to determine significance (ns: non-significant). Horizontal lines show the mean and error bars represent the standard deviation. The numbers of data points obtained from experiments are indicated on the graphs in parentheses.

Our data aligns with some previous reports while partially contradicting some others. For example, *Rubio-Sánchez et al* showed size-dependent partitioning of oligonucleotides in model membranes where larger oligos partitioned less into the ordered domains which is in line with our observations with proteins (27). However, by using detergent resistance assay, *Shao et al* showed that O-glycans direct some adhesion molecules to ordered membrane domains which contrasts with our findings (53). This discrepancy might be due to the different experimental systems (detergent resistance vs. model membranes) or investigated glycosylation patterns (O-glycosylation vs. overall glycosylation).

While our data could show that glycosylation and the apparent molecular mass (as we summarized with the term “size”) are determinants of partitioning, the exact physical mechanism is not trivial to unravel since we do not have sufficient information on the exact geometry of the reconstituted proteins on membranes such as rigidity, bulkiness, length, apparent radius and bending. Therefore, we cannot use such parameters in a quantitative model. However, a plausible speculation would be the excluded volume interactions: larger, glycosylated, and charged (due to sialic acid) molecules might repulse each other from tightly packed ordered domains (since lipid per area, hence molecular proximity, is higher in L_o_ phase) which might account for their exclusion from ordered domains. This effect can be attributed to apparent molecular mass, but also to other properties such as charge, rigidity, bulkiness, length, or apparent radius of the molecule. With our current knowledge, we have the exact information about the apparent molecular mass of the proteins, which is perfectly in trend with observed partitioning for the proteins we used. However, the other factors might better account for the observed effects, which need to be addressed in the future with more experimental and in silico work.

### Apparent ECD mass influences molecular diffusion

Another dynamic behavior of proteins is their diffusion as it determines their interaction dynamics with other macromolecules. There is still a controversy about the structural determinants of membrane protein mobility. From the very first diffusion model (35) to the succeeding modifications (38–42), none of them explicitly takes ECD size into account. There is a consensus in the field that protein diffusion in the membrane is determined by the TMD, the lipid environment, and their attachment to the membrane. To investigate the effect of ECDs on protein dynamics in the membrane, we measured the diffusion coefficients of proteins using FCS, where fluorescence intensity fluctuations were used to obtain information about molecular diffusion **(Fig. 4A)**. We first tested how the diffusion of ECDs was affected by their apparent mass in GUVs. For this purpose, we measured the diffusion of ECDs in POPC doped with 1 mol% DO Ni-NTA, and we observed a reciprocal relationship between the mobility and the apparent ECD mass, that is, an increase in the size of the protein leads to slower diffusion **(Fig. 4B)**. The same relationship was observed in GUVs of different lipid compositions as well **(Supp. Fig. 5)**. The diffusion of the lipid analogue AbStR-DPPE displays no change in GUVs with ECDs of different sizes **(Supp. Fig. 6)**, confirming that the membrane physical parameters (e.g., fluidity) and the experimental conditions (e.g., temperature fluctuations), did not cause the differences in diffusion for different ECDs. Moreover, the number of particles in each experimental set-up for all proteins was comparable, ruling out the effect of crowding (46) on observed differences in diffusion **(Supp. Fig. 7)**. To evaluate the role of glycosylation and apparent vs. predicted mass on diffusion, as we did for partitioning, we used CD34, PODXL, and PODXL2. Similar to partitioning, we observed a correlation between diffusion and apparent mass but not with predicted mass **(Figure 4C, D, Supp. Fig. 8-10, Supp. Table 2, 3)**. When we treated these proteins with deglycosylation enzymes, PODXL2 and PODXL sped up significantly while CD34 did not change notably, similar to partitioning experiments **(Figure 4E, Supp. Fig. 11)**.

**Fig. 4.**
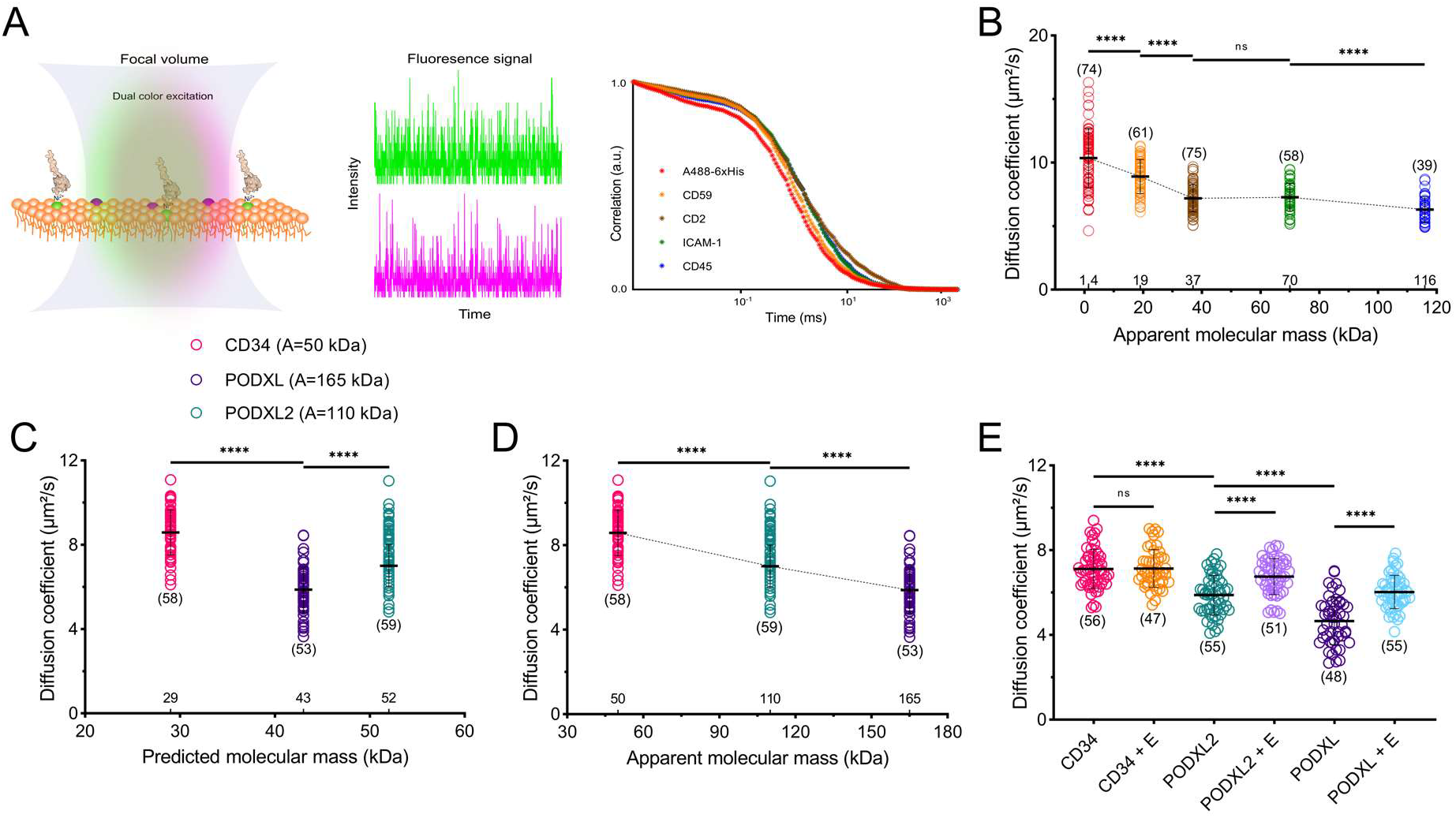
Glycosylation and apparent ECD mass determine the protein diffusion in GUVs. (A) Illustration of the FCS experiment. (B) Diffusion coefficients of the different ECDs in GUVs consisting of POPC. Student’s t-test (two-tailed, non-parameterized) was used to determine significance (****P<0.0001; ns: non-significant). (C) Diffusion coefficients of CD34, PODXL and PODXL2 as a function of the predicted mass and (D) as a function of the apparent mass. (E) Diffusion coefficients of CD34, PODXL and PODXL2 after deglycosylation enzyme treatment. Horizontal lines show the mean and error bars represent the standard deviation. The numbers of data points obtained from experiments are indicated on the graphs in parentheses.

Our findings support some of the recent reports (45, 47, 48) while partially contradicting some others (21, 46). For example, *Houser et al* found size-dependent diffusion only in crowded conditions while we observed this phenomenon in non-crowded conditions, as well (46). The likely cause of this could be a different model membrane system (supported bilayers vs. free-standing membranes). Moreover, *Popik et al* did not find glycosylation as a factor changing the ordered domain localization while our data shows a clear dependence of partitioning on the glycosylation of the proteins (21). This could be due to discrepancies of experimental modalities (e.g., detergent resistance assay vs. model membranes). Our results are complementary to those of *Hartel et al*., as they investigate the impact of size extension/truncation of the pathogen GPI-anchored VSG glycoprotein (45). They reveal a correlation between diffusion and the ectodomain size which can be different for different variants of the protein. With our study, we can extend the finding of size-dependent mobility to PM residing proteins, in a more inclusive approach.

### The GPMV membrane environment reduces the size effect on molecular mobility

The cellular PM is a more crowded and complex environment than GUVs. The size effect on diffusion can be enhanced or diluted in crowded environments. For example, *Houser et al* showed that in crowded environments, the size of the proteins plays a greater role in their diffusion compared to non-crowded environment (46). To test how our proteins behave in near-physiological crowding, we performed diffusion experiments on GPMVs **(Fig. 5A)**. The variation in membrane compositions between individual GPMVs leads to larger variability of the diffusion coefficient, therefore the data spread in graphs is larger **(Fig. 5B)**. Despite the large standard deviations, diffusion of proteins in GPMVs generally resembles the trend in GUVs with certain exceptions. The diffusion coefficient decreases with an increase in ECD size for the first three small molecules. For relatively larger proteins, diffusion did not differ drastically. AbStR-DPPE shows no significant change in diffusion with ECDs of different sizes, confirming that the observed differences are indeed due to ECD size **(Supp. Fig. 12)**. This data means that in GPMVs, the size of ECDs is important, but in contrast to GUVs, small differences in size can be masked by other factors in the native membranes such as molecular crowding. However, additional factors in GPMVs might be responsible for this effect. For example, GPMVs exhibit negatively charged lipids in the outer leaflet which ECDs can interact with. In contrast, in live cells (or in GUVs), there are not negatively charged lipids exposed in the outer leaflet. Second, chemicals used for vesiculation can lead to unforeseen effects on protein mobility. Moreover, incorporation of nickel-chelating lipids into GPMVs is a burdensome protocol which might change the GPMV membranes. Therefore, the results obtained from GUVs and GPMVs cannot be compared directly in the context of crowding. Yet, our results display an indication of possible effects of size in two different model membrane systems of varying membrane complexity.

**Fig. 5.**
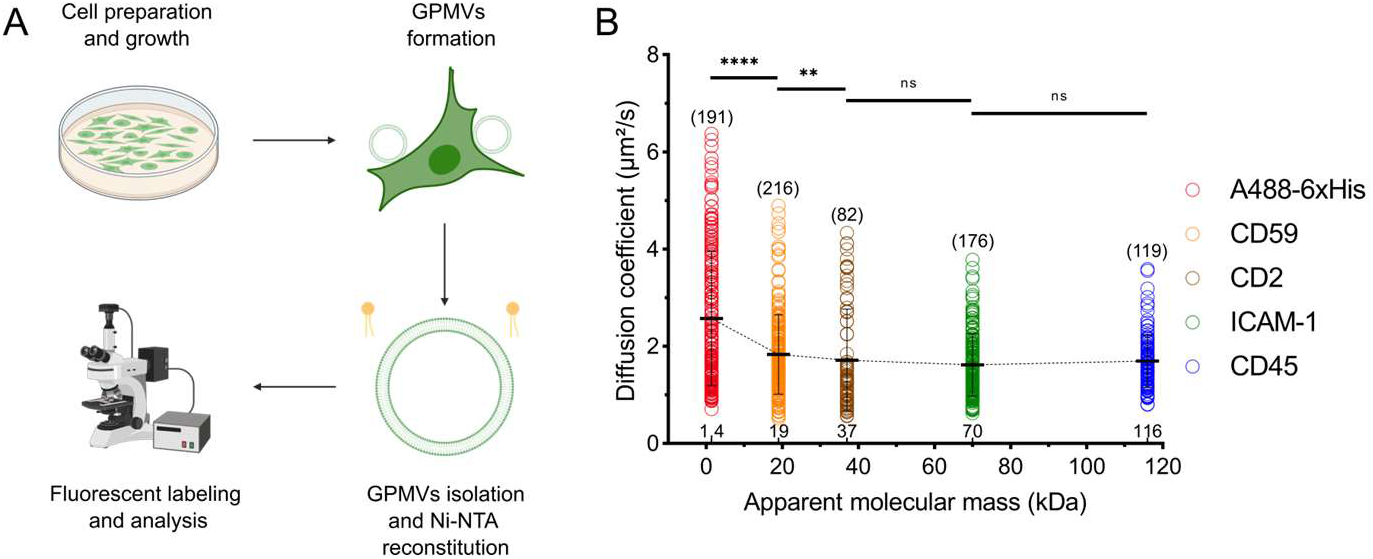
ECD size and protein diffusion in GPMVs. (A) Preparation of GPMVs functionalized with Ni-NTA (B) Diffusion of the different ECDs in GPMVs. Student’s t-test (two-tailed, non-parameterized) was used to determine significance (****P<0.0001; **P=0.0054; ns: non-significant). Horizontal lines show the mean and error bars represent the standard deviation. The numbers of data points obtained from experiments are indicated on the graphs in parentheses.

## CONCLUSION

In this study, we investigated the largely overlooked role of ECD size on MP dynamics. Our results show a decrease both in partitioning into ordered domains and in diffusion coefficient with the increase in ECD size. In the light of our data, we suggest that ECD is a necessary component for understanding protein behavior in the PM. Our data highlights the necessity of new models explaining the diffusion and partitioning of proteins taking their extracellular size into account. Accumulating experimental and in silico data will eventually lead to more refined and accurate models.

Our data has potential implications for the fundamental understanding of cellular biology. Differences in ECD size, for example, between isotypes or caused by post-translational modifications, ligand binding, or enzymatic cleavage could lead to different protein mobility and localization in the PM, which could be required for certain functions. Therefore, ECD size might serve as a parameter for the compartmentalization of membrane components by influencing their mobility and domain partitioning. This postulation is in line with recent evidence on how glycosylation influences the protein compartmentalization and cellular signaling (53, 56, 57). In this regard, our data can be the connection between different mechanisms proposed for membrane protein reorganization during signaling initiation since it shows the correlation between ECD size, diffusion, and domain partitioning.

There are currently multiple mechanisms proposed for T cell signaling (and most are applicable to other immune cell types). Size-dependent, lipid-dependent, or diffusion-dependent exclusion of inhibitory molecules (such as CD45) from the immune synapse are three of the main and seemingly distant mechanisms explaining T-cell signaling (3–8). However, our results suggest that all three phenomena can be intertwined. The extracellular size, lipid domain partitioning, and molecular diffusion are not entirely independent. Tall and heavily glycosylated (hence more rigid) molecules can be driven out of a certain lipid environment and diffuse slower due to their size. Of course, the size-dependent partitioning is unlikely to be the only mechanism determining whether immune cells will be activated or not; however, it would be a plausible fine-tuning mechanism. The exact role of size-dependent partitioning of proteins in immune signaling needs to be tested in the future. Similar to size-dependent partitioning, size-dependent diffusion of molecules can also fine-tune signaling output by determining the collision rate between the receptor, the activating, and inhibitory proteins. The activating molecules for immune signaling (e.g., Lck kinase that phosphorylates the T-cell receptors) are usually small proteins while inhibitory molecules (e.g., CD45 phosphatase that dephosphorylates the T-cell receptors) bear large ECDs. This discrepancy likely plays a role in their diffusion. Since diffusion rate is the currency for molecular interactions, size-dependent diffusion is a potential effector of immune signaling.

It is, however, essential to acknowledge the limitations of our study. The direct translatability from model membrane systems to the native PM requires caution. The absence of other membrane proteins (i.e., absence of protein-protein interactions), the cortical actin cytoskeleton, and the extracellular networks (i.e., lack of crowding, ECM-protein interactions(34)) might hinder the direct translatability of GUV experiments. In addition, all ECDs but CD59 originate from integral proteins, hence, we refrain from drawing conclusions on the full-length proteins. These questions can be studied in GPMVs bearing the full-length proteins with the same TMDs but varying ECDs to be able to investigate these phenomena in a more physiological setting. However, GPMVs also have several shortcomings, such as the lack of compositional control compared to GUVs, and unpredicted effects of chemicals used for GPMV formation. Another possible caveat is the unpredictable geometry of ECDs (e.g., bending towards the membrane) due to the nature of the chemical coupling we used. However, previous studies confirmed that the height of the molecules is reasonably preserved when they are coupled to the membranes via His-Ni-NTA chemistry (58). Due to the sole focus on the ECD fraction of the proteins via membrane anchoring in this study, future validation experiments investigating the impact of the ECD in transmembrane proteins will strengthen our findings.

Despite the aforementioned limitations, our system provides a clean and controllable environment to evaluate how ECD size influences the dynamic behavior of the proteins independent of their TMDs or ICDs. We believe that this work will pave the way for further experiments and simulations, which will eventually lead to new models for protein dynamics in the PM.

## ACKNOWLEDGMENTS

We thank Hannes Feyrer and CSI:Nano Lab for their feedback on figures and Dr. Markus Deserno for discussing our results. SciLifeLab Advanced Light Microscopy facility and National Microscopy Infrastructure (VR-RFI 2016-00968) for their support on imaging. COG was supported by a SciLifeLab summer internship (C-2021-0466), ES is supported by grants from Swedish Research Council Starting Grant (2020-02682), from Karolinska Institutet and from the SciLifeLab National COVID-19 Research Program, financed by the Knut and Alice Wallenberg Foundation. Alphafold model analysis was performed via USCS ChimeraX developed by the Resource for Biocomputing, Visualization and Informatics at the University of California, San Francisco, with support from National Institutes of Health R01-GM129325 and the Office of Cyber Infrastructure and Computational Biology, National Institute of Allergy and Infectious Diseases. The figures were prepared by Inkscape and BioRender.

## REFERENCES

1. Fagerberg, L., K. Jonasson, G. von Heijne, M. Uhlén, and L. Berglund. 2010. Prediction of the human membrane proteome. Proteomics. 10:1141–1149.

2. Simons, K., and D. Toomre. 2000. Lipid rafts and signal transduction. Nat Rev Mol Cell Biol. 1:31–39.

3. Davis, S.J., and P.A. van der Merwe. 2006. The kinetic-segregation model: TCR triggering and beyond. Nature immunology. 7:803–9.

4. Felce, J., E. Sezgin, M. Wane, H. Brouwer, M.L. Dustin, C. Eggeling, and S.J. Davis. 2018. CD45 exclusion and cross-linking based receptor signaling together broaden FcεRI reactivity. Science Signalling. 11:eaat0756.

5. Bakalar, M.H., A.M. Joffe, E.M. Schmid, S. Son, M. Podolski, and D.A. Fletcher. 2018. Size-Dependent Segregation Controls Macrophage Phagocytosis of Antibody-Opsonized Targets. Cell. 174:131-142.e13.

6. Swamy, M., K. Beck-Garcia, E. Beck-Garcia, F.A. Hartl, A. Morath, O.S. Yousefi, E.P. Dopfer, E. Molnár, A.K. Schulze, R. Blanco, A. Borroto, N. Martín-Blanco, B. Alarcon, T. Höfer, S. Minguet, and W.W.A. Schamel. 2016. A Cholesterol-Based Allostery Model of T Cell Receptor Phosphorylation. Immunity. 44:1091–1101.

7. Chen, K.Y., E. Jenkins, M. Körbel, A. Ponjavic, A.H. Lippert, A.M. Santos, N. Ashman, C. O’Brien-Ball, J. McBride, D. Klenerman, and S.J. Davis. 2021. Trapping or slowing the diffusion of T cell receptors at close contacts initiates T cell signaling. Proc Natl Acad Sci U S A. 118:e2024250118.

8. Urbančič, I., L. Schiffelers, E. Jenkins, W. Gong, A.M. Santos, F. Schneider, C. O’Brien-Ball, M.T. Vuong, N. Ashman, E. Sezgin, and C. Eggeling. 2021. Aggregation and mobility of membrane proteins interplay with local lipid order in the plasma membrane of T cells. FEBS Lett. 595:2127–2146.

9. Sych, T., C.O. Gurdap, L. Wedemann, and E. Sezgin. 2021. How Does Liquid-Liquid Phase Separation in Model Membranes Reflect Cell Membrane Heterogeneity? Membranes. 11:323.

10. Sezgin, E., I. Levental, M. Grzybek, G. Schwarzmann, V. Mueller, A. Honigmann, V.N. Belov, C. Eggeling, Ü. Coskun, K. Simons, and P. Schwille. 2012. Partitioning, diffusion, and ligand binding of raft lipid analogs in model and cellular plasma membranes. Biochimica et Biophysica Acta (BBA) - Biomembranes. 1818:1777–1784.

11. Lorent, J.H., and I. Levental. 2015. Structural determinants of protein partitioning into ordered membrane domains and lipid rafts. Chemistry and Physics of Lipids. 192:23–32.

12. Polley, A., A. Orlowski, R. Danne, A.A. Gurtovenko, J. Bernardino de la Serna, C. Eggeling, S.J. Davis, T. Rog, and I. Vattulainen. 2017. Glycosylation and Lipids Working in Concert Direct CD2 Ectodomain Orientation and Presentation. The journal of physical chemistry letters. 1060–1066.

13. Stone, M.B., S.A. Shelby, M.F. Núñez, K. Wisser, and S.L. Veatch. 2017. Protein sorting by lipid phase-like domains supports emergent signaling function in B lymphocyte plasma membranes. eLife. 6:e19891.

14. Yang, S.-T., A.J.B. Kreutzberger, V. Kiessling, B.K. Ganser-Pornillos, J.M. White, and L.K. Tamm. 2017. HIV virions sense plasma membrane heterogeneity for cell entry. Sci Adv. 3:e1700338.

15. Sharpe, H.J., T.J. Stevens, and S. Munro. 2010. A Comprehensive Comparison of Transmembrane Domains Reveals Organelle-Specific Properties. Cell. 142:158–169.

16. Diaz-Rohrer, B.B., K.R. Levental, K. Simons, and I. Levental. 2014. Membrane raft association is a determinant of plasma membrane localization. PNAS. 111:8500–8505.

17. Lorent, J.H., B. Diaz-Rohrer, X. Lin, K. Spring, A.A. Gorfe, K.R. Levental, and I. Levental. 2017. Structural determinants and functional consequences of protein affinity for membrane rafts. Nat Commun. 8:1219.

18. Nalivaeva, N.N., and A.J. Turner. 2009. Lipid Anchors to Proteins. In: Lajtha A, G Tettamanti, G Goracci, editors. Handbook of Neurochemistry and Molecular Neurobiology: Neural Lipids. Boston, MA: Springer US. pp. 353–372.

19. van Deventer, S., A.B. Arp, and A.B. van Spriel. 2021. Dynamic Plasma Membrane Organization: A Complex Symphony. Trends Cell Biol. 31:119–129.

20. Sezgin, E., I. Levental, S. Mayor, and C. Eggeling. 2017. The mystery of membrane organization: composition, regulation and physiological relevance of lipid rafts. Nat Rev Mol Cell Biol. 18:361–374.

21. Popik, W., and T.M. Alce. 2004. CD4 Receptor Localized to Non-raft Membrane Microdomains Supports HIV-1 Entry: IDENTIFICATION OF A NOVEL RAFT LOCALIZATION MARKER IN CD4 *. Journal of Biological Chemistry. 279:704–712.

22. Stachowiak, J.C., E.M. Schmid, C.J. Ryan, H.S. Ann, D.Y. Sasaki, M.B. Sherman, P.L. Geissler, D.A. Fletcher, and C.C. Hayden. 2012. Membrane bending by protein–protein crowding. Nat Cell Biol. 14:944–949.

23. Yuan, F., H. Alimohamadi, B. Bakka, A.N. Trementozzi, K.J. Day, N.L. Fawzi, P. Rangamani, and J.C. Stachowiak. 2021. Membrane bending by protein phase separation. PNAS. 118.

24. Banjade, S., and M.K. Rosen. 2014. Phase transitions of multivalent proteins can promote clustering of membrane receptors. eLife. 3:e04123.

25. Zeno, W.F., K.E. Johnson, D.Y. Sasaki, S.H. Risbud, and M.L. Longo. 2016. Dynamics of Crowding-Induced Mixing in Phase Separated Lipid Bilayers. J Phys Chem B. 120:11180–11190.

26. Varshney, P., V. Yadav, and N. Saini. 2016. Lipid rafts in immune signalling: current progress and future perspective. Immunology. 149:13–24.

27. Rubio-Sánchez, R., S.E. Barker, M. Walczak, P. Cicuta, and L.D. Michele. 2021. A Modular, Dynamic, DNA-Based Platform for Regulating Cargo Distribution and Transport between Lipid Domains. Nano Lett. 21:2800–2808.

28. Fujiwara, T., K. Ritchie, H. Murakoshi, K. Jacobson, and A. Kusumi. 2002. Phospholipids undergo hop diffusion in compartmentalized cell membrane. Journal of Cell Biology. 157:1071–1081.

29. Wawrezinieck, L., H. Rigneault, D. Marguet, and P.F. Lenne. 2005. Fluorescence correlation spectroscopy diffusion laws to probe the submicron cell membrane organization. Biophysical journal. 89:4029–42.

30. Lenne, P.F., L. Wawrezinieck, F. Conchonaud, O. Wurtz, A. Boned, X.J. Guo, H. Rigneault, H.T. He, and D. Marguet. 2006. Dynamic molecular confinement in the plasma membrane by microdomains and the cytoskeleton meshwork. The EMBO journal. 25:3245–56.

31. Eggeling, C., C. Ringemann, R. Medda, G. Schwarzmann, K. Sandhoff, S. Polyakova, V.N. Belov, B. Hein, C. von Middendorff, A. Schonle, and S.W. Hell. 2009. Direct observation of the nanoscale dynamics of membrane lipids in a living cell. Nature. 457:1159–62.

32. Alenghat, F.J., and D.E. Golan. 2013. Membrane protein dynamics and functional implications in mammalian cells. Curr Top Membr. 72:89–120.

33. Schneider, F., D. Waithe, M.P. Clausen, S. Galiani, T. Koller, G. Ozhan, C. Eggeling, and E. Sezgin. 2017. Diffusion of lipids and GPI-anchored proteins in actin-free plasma membrane vesicles measured by STED-FCS. Mol Biol Cell. 28:1507–1518.

34. Frischknecht, R., M. Heine, D. Perrais, C.I. Seidenbecher, D. Choquet, and E.D. Gundelfinger. 2009. Brain extracellular matrix affects AMPA receptor lateral mobility and short-term synaptic plasticity. Nat Neurosci. 12:897–904.

35. Saffman, P.G., and M. Delbrück. 1975. Brownian motion in biological membranes. PNAS. 72:3111–3113.

36. Hughes, B.D., B.A. Pailthorpe, and L.R. White. 1981. The translational and rotational drag on a cylinder moving in a membrane. Journal of Fluid Mechanics. 110:349–372.

37. Guigas, G., and M. Weiss. 2006. Size-dependent diffusion of membrane inclusions. Biophys J. 91:2393–2398.

38. Frick, M., K. Schmidt, and B.J. Nichols. 2007. Modulation of Lateral Diffusion in the Plasma Membrane by Protein Density. Current Biology. 17:462–467.

39. Petrov, E.P., and P. Schwille. 2008. Translational Diffusion in Lipid Membranes beyond the Saffman-Delbrück Approximation. Biophysical Journal. 94:L41–L43.

40. Ramadurai, S., A. Holt, V. Krasnikov, G. van den Bogaart, J.A. Killian, and B. Poolman. 2009. Lateral Diffusion of Membrane Proteins. J. Am. Chem. Soc. 131:12650–12656.

41. Morozova, D., G. Guigas, and M. Weiss. 2011. Dynamic Structure Formation of Peripheral Membrane Proteins. PLOS Computational Biology. 7:e1002067.

42. Goose, J.E., and M.S.P. Sansom. 2013. Reduced Lateral Mobility of Lipids and Proteins in Crowded Membranes. PLOS Computational Biology. 9:e1003033.

43. Weiß, K., A. Neef, Q. Van, S. Kramer, I. Gregor, and J. Enderlein. 2013. Quantifying the Diffusion of Membrane Proteins and Peptides in Black Lipid Membranes with 2-Focus Fluorescence Correlation Spectroscopy. Biophys J. 105:455–462.

44. Wieser, S., M. Moertelmaier, E. Fuertbauer, H. Stockinger, and G.J. Schütz. 2007. (Un)Confined Diffusion of CD59 in the Plasma Membrane Determined by High-Resolution Single Molecule Microscopy. Biophysical Journal. 92:3719–3728.

45. Hartel, A.J.W., M. Glogger, G. Guigas, N.G. Jones, S.F. Fenz, M. Weiss, and M. Engstler. 2015. The molecular size of the extra-membrane domain influences the diffusion of the GPI-anchored VSG on the trypanosome plasma membrane. Sci Rep. 5:10394.

46. Houser, J.R., D.J. Busch, D.R. Bell, B. Li, P. Ren, and J.C. Stachowiak. 2016. The impact of physiological crowding on the diffusivity of membrane bound proteins. Soft Matter. 12:2127–2134.

47. Guigas, G., and M. Weiss. 2015. Membrane protein mobility depends on the length of extra-membrane domains and on the protein concentration. Soft Matter. 11:33–37.

48. Kim, D.-H., K. Zhou, D.-K. Kim, S. Park, J. Noh, Y. Kwon, D. Kim, N.W. Song, J.-B. Lee, P.-G. Suh, N.K. Lee, and S.H. Ryu. 2015. Analysis of Interactions between the Epidermal Growth Factor Receptor and Soluble Ligands on the Basis of Single-Molecule Diffusivity in the Membrane of Living Cells. Angewandte Chemie. 127:7134–7138.

49. Schindelin, J., I. Arganda-Carreras, E. Frise, V. Kaynig, M. Longair, T. Pietzsch, S. Preibisch, C. Rueden, S. Saalfeld, B. Schmid, J.Y. Tinevez, D.J. White, V. Hartenstein, K. Eliceiri, P. Tomancak, and A. Cardona. 2012. Fiji: an open-source platform for biological-image analysis. Nature methods. 9:676–82.

50. Waithe, D., M.P. Clausen, E. Sezgin, and C. Eggeling. 2016. FoCuS-point: software for STED fluorescence correlation and time-gated single photon counting. Bioinformatics. 32:958–960.

51. Yang, H., and E.L. Reinherz. 2001. Dynamic recruitment of human CD2 into lipid rafts. Linkage to T cell signal transduction. J Biol Chem. 276:18775–18785.

52. Yamabhai, M., and R.G.W. Anderson. 2002. Second cysteine-rich region of epidermal growth factor receptor contains targeting information for caveolae/rafts. J Biol Chem. 277:24843–24846.

53. Shao, B., T. Yago, H. Setiadi, Y. Wang, P. Mehta-D’souza, J. Fu, P.R. Crocker, W. Rodgers, L. Xia, and R.P. McEver. 2015. O-glycans direct selectin ligands to lipid rafts on leukocytes. Proceedings of the National Academy of Sciences. 112:8661–8666.

54. Wang, L., J. Zhang, D. Wang, and C. Song. 2022. Membrane contact probability: An essential and predictive character for the structural and functional studies of membrane proteins. PLOS Computational Biology. 18:e1009972.

55. Lu, C.-H., K. Pedram, C.-T. Tsai, T. Jones, X. Li, M.L. Nakamoto, C.R. Bertozzi, and B. Cui. 2022. Membrane curvature regulates the spatial distribution of bulky glycoproteins. Nat Commun. 13:3093.

56. Wasim, L., F.H.M. Buhari, M. Yoganathan, T. Sicard, J. Ereño-Orbea, J.-P. Julien, and B. Treanor. 2019. N-Linked Glycosylation Regulates CD22 Organization and Function. Frontiers in Immunology. 10.

57. Blouin, C.M., Y. Hamon, P. Gonnord, C. Boularan, J. Kagan, C. Viaris de Lesegno, R. Ruez, S. Mailfert, N. Bertaux, D. Loew, C. Wunder, L. Johannes, G. Vogt, F.X. Contreras, D. Marguet, J.L. Casanova, C. Gales, H.T. He, and C. Lamaze. 2016. Glycosylation-Dependent IFN-gammaR Partitioning in Lipid and Actin Nanodomains Is Critical for JAK Activation. Cell. 166:920–34.

58. Son, S., S.C. Takatori, B. Belardi, M. Podolski, M.H. Bakalar, and D.A. Fletcher. 2020. Molecular height measurement by cell surface optical profilometry (CSOP). Proc. Natl. Acad. Sci. U.S.A. 117:14209–14219.

