## Supplementary Information for "Influence of the extracellular domain size on the dynamic behavior of membrane proteins"

**Supplementary Data**

**Supplementary Figures**

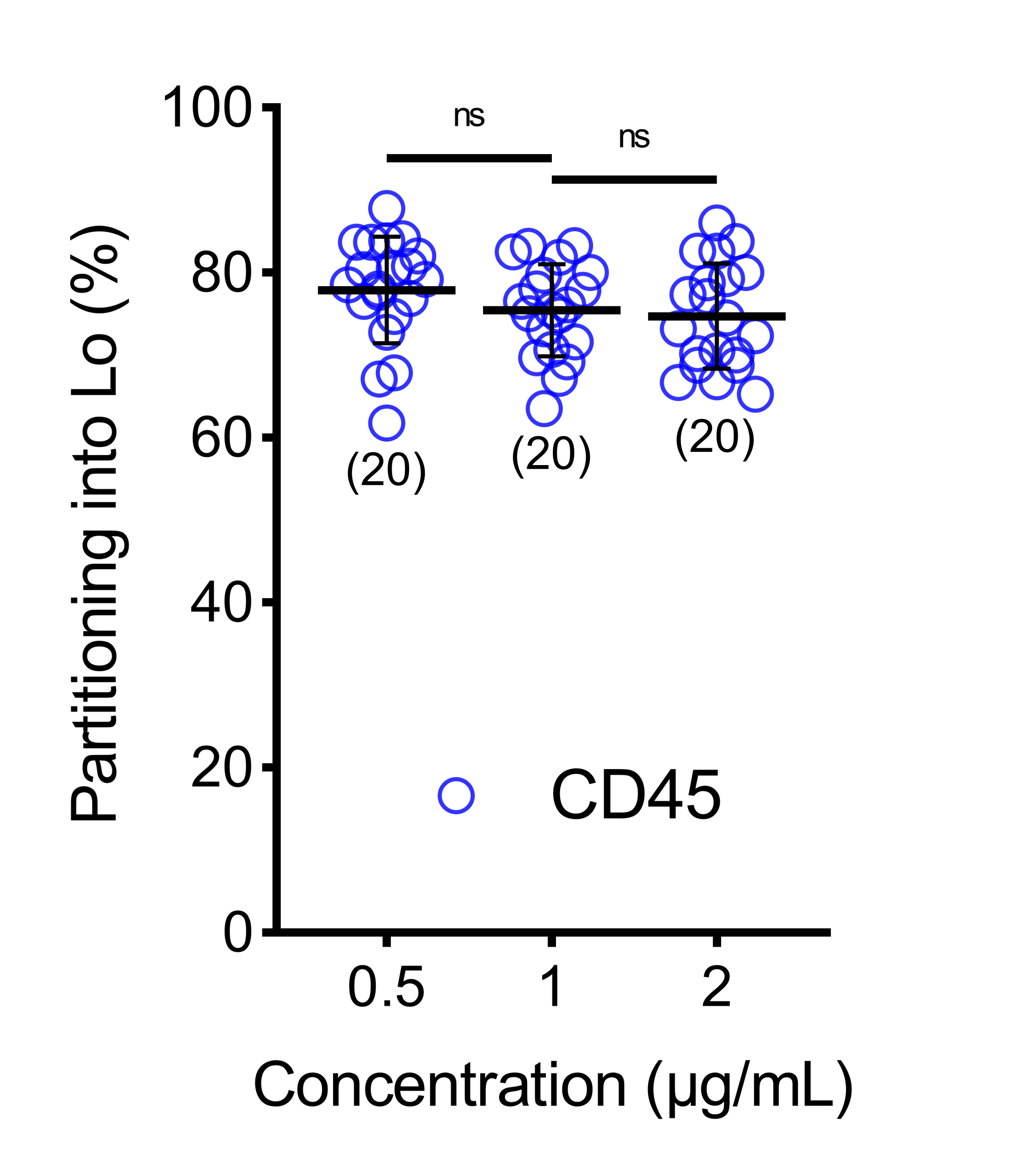

**Supp. Fig. 1. The partitioning of CD45 (the largest protein) remains constant across varying protein concentrations.** A non-parameterized ANOVA was used to determine significance (ns: non-significant). Horizontal lines show the mean and error bars represent the standard deviation. The number of data points obtained for each experiment are indicated on the graphs in parentheses.

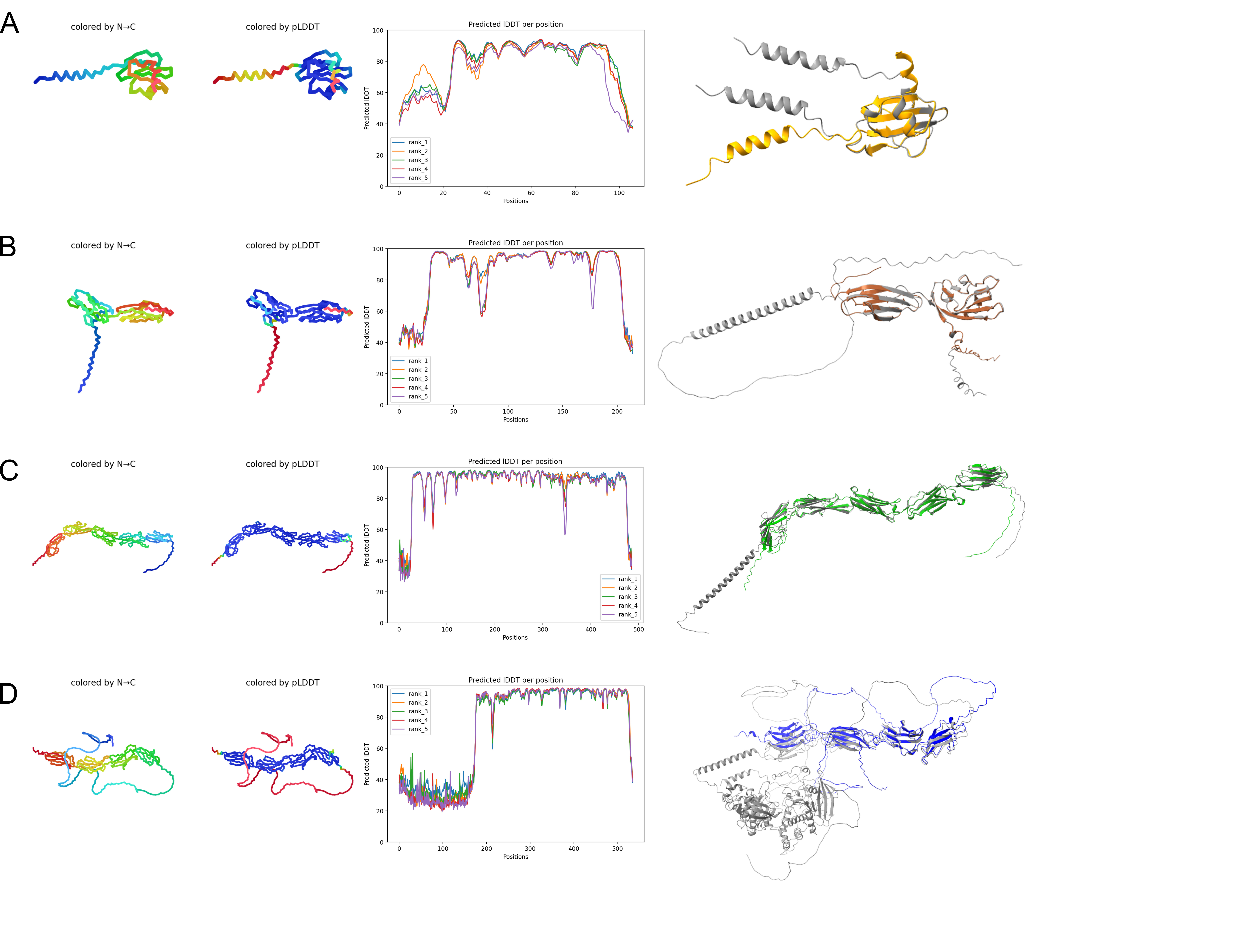
**Supp. Fig. 2. ECD Structural models via ALPHAFOLD2.** Model obtained via Collabfold, multiple sequence alignment performed via Jackhmmer, displayed is the highest ranked out of 5 models performed, with the LDDT graph for all models displayed next to the structures. Also shown are alignments to published structures for the whole protein from PDB, each being alphafold models, PDB code indicated with the protein as following: (A) CD59, AlphaFold: AF-P13987-F1, (B) CD2, AlphaFold: AF_P06729-F1, (C) ICAM-1, AlphaFold: AF-P05362-F1, (D) CD45, AlphaFold: AF-P08575-F1. Some of the structures cannot be resolved with Alphafold, presumably due to glycosylation.

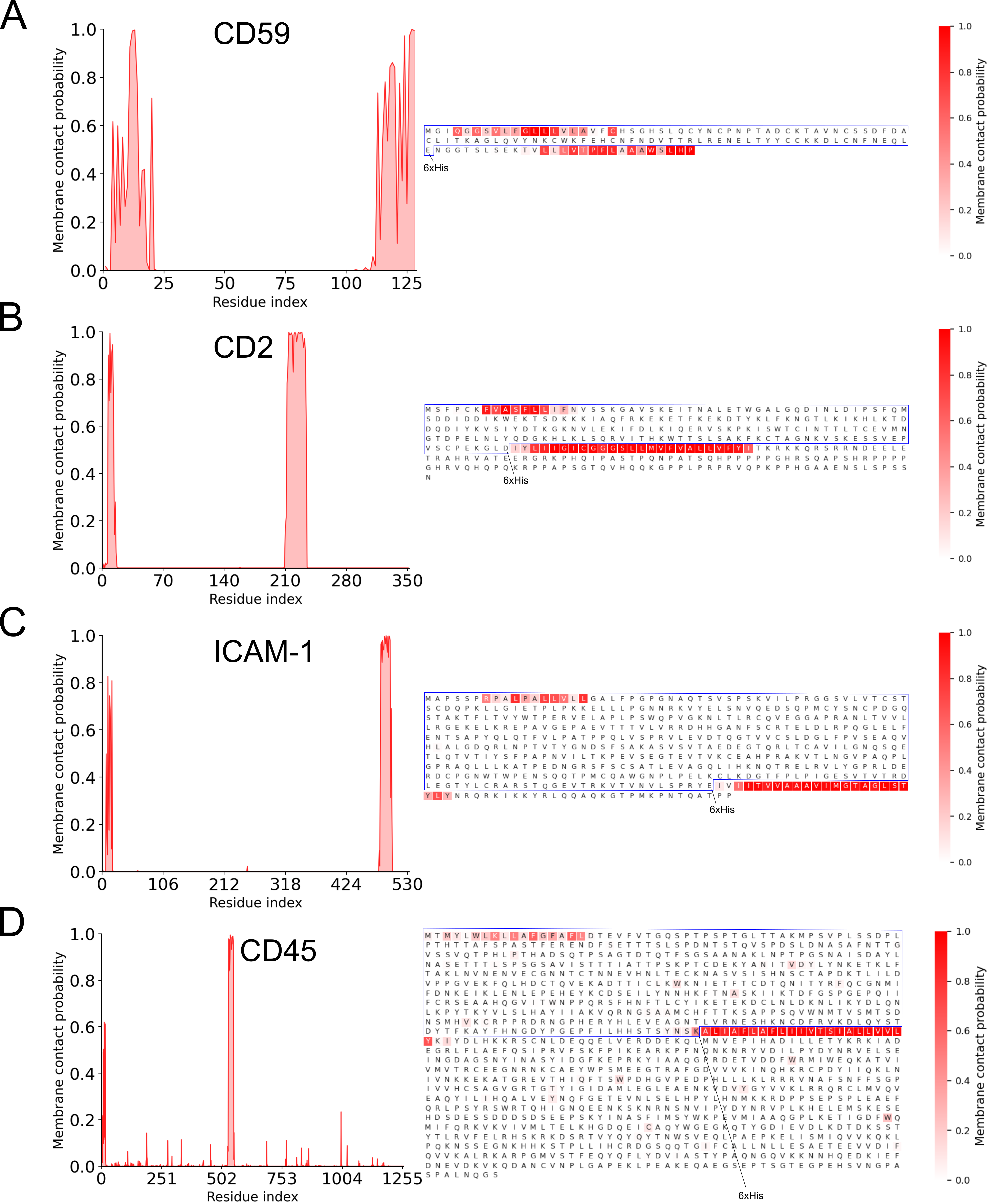

**Supp. Fig. 3. Membrane contact probability.** The analysis was done on full-length proteins and blue frames show extracellular domain of the proteins used in the study. 6xHis-tag in C-terminus is shown with arrow for each protein. Amino acids next the membrane anchoring sites can result in protein lipid interactions. Each protein has similar hydrophobic residues towards the far end of the proteins.

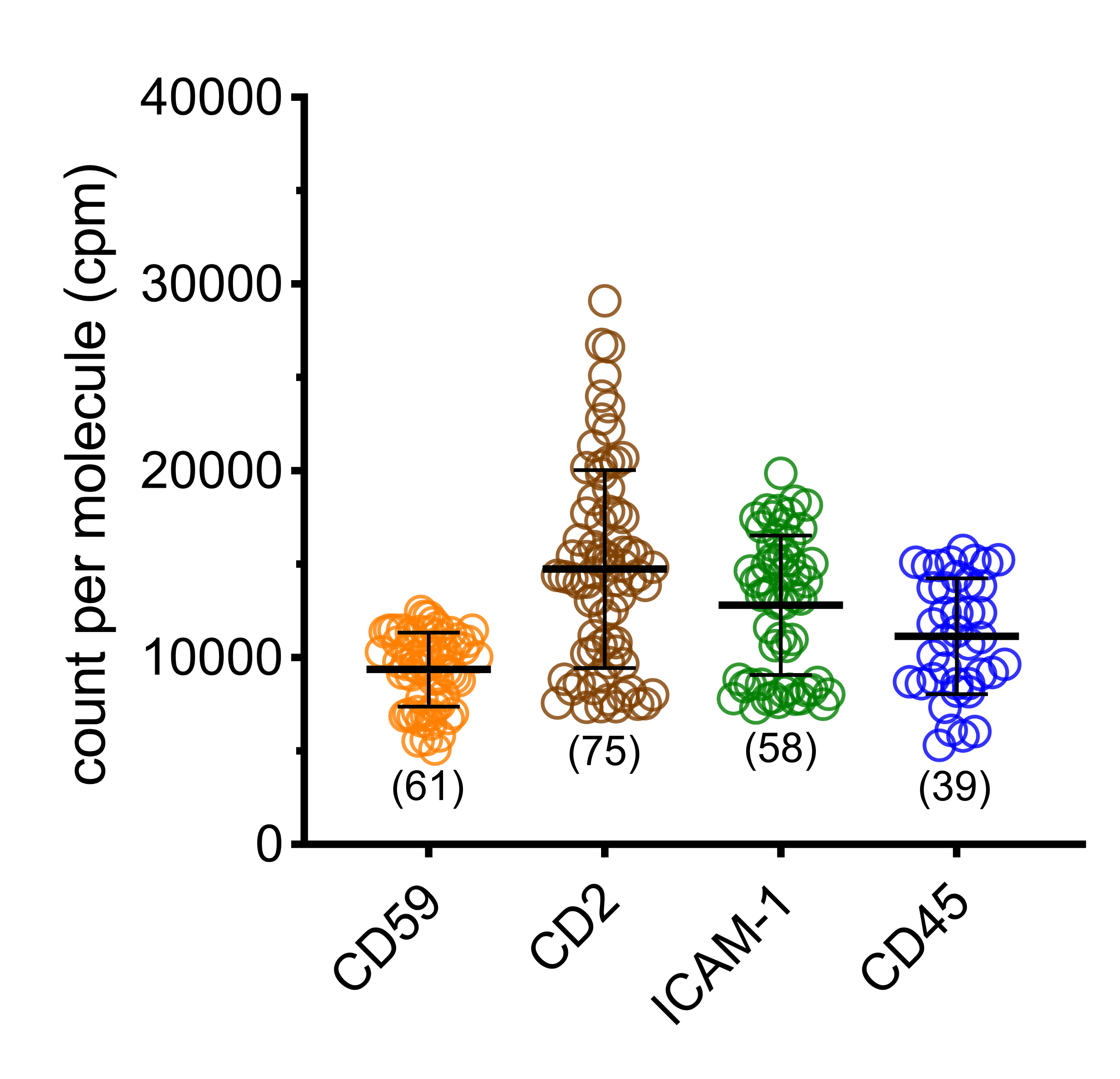

**Supp. Fig. 4. Count per molecule for FCS measurement on GUVs.** Horizontal lines show the mean and error bars represent the standard deviation. The number of data points obtained from experiments are indicated on the graphs in parentheses. Data does not show a trend that can correlate with diffusion or partitioning.

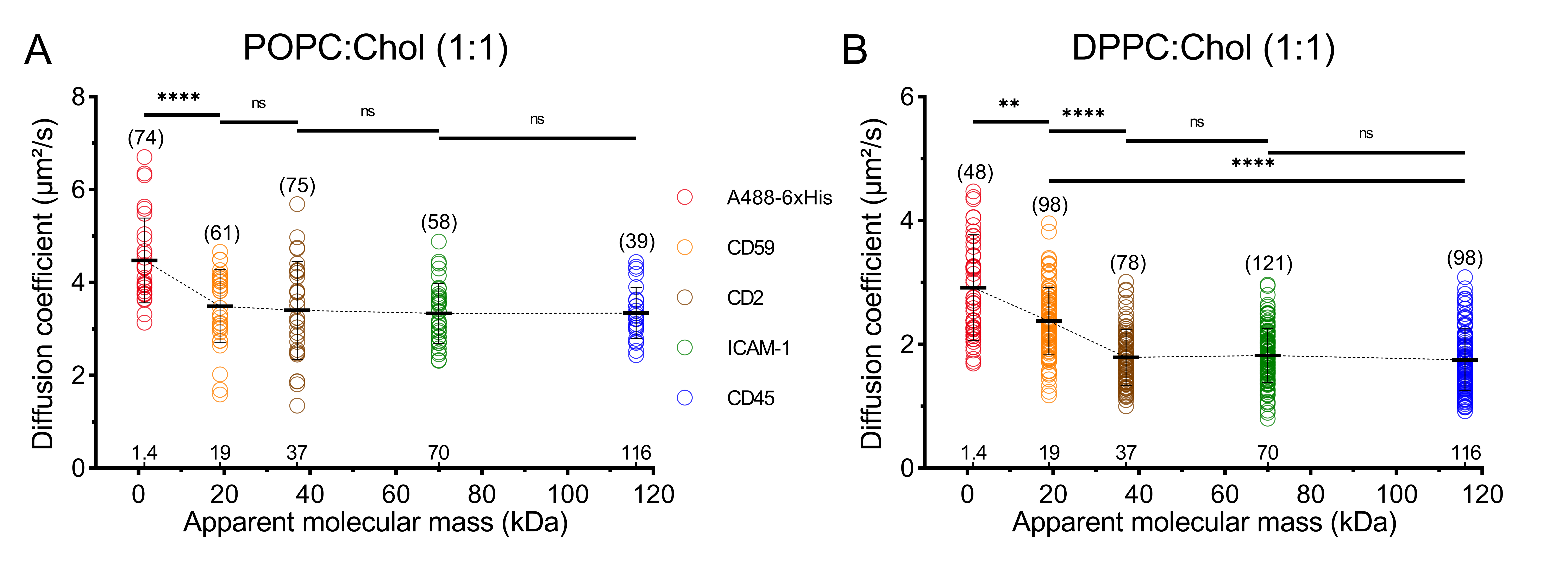

**Supp. Fig. 5. Diffusion coefficients of the different ECDs in GUVs with (A) POPC:Chol (1:1) and (B) with DPPC/Chol (1:1).** Student’s t-test (two-tailed, non-parameterized) was used to determine significance (****P<0.0001; **P=0.001; ns: non-significant). Horizontal lines show the mean and error bars represent the standard deviation. The number of data points obtained from experiments are indicated on the graphs in parentheses.

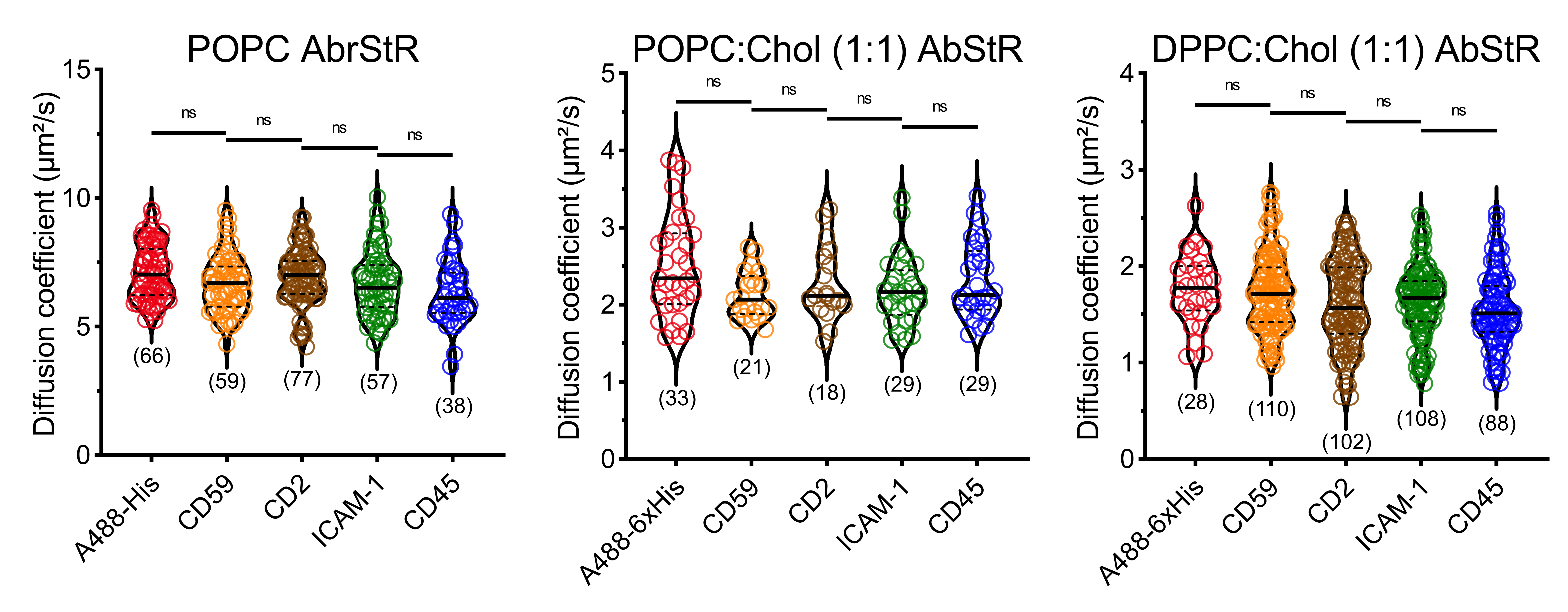

**Supp. Fig. 6. Diffusion of membrane dye (AbStR-DPPE) in GUVs.** Non-parameterized ANOVA was used to determine significance (ns: non-significant). Horizontal thick lines show the median and dotted lines represent the quartiles. The number of data points obtained from experiments are indicated on the graphs in parentheses.

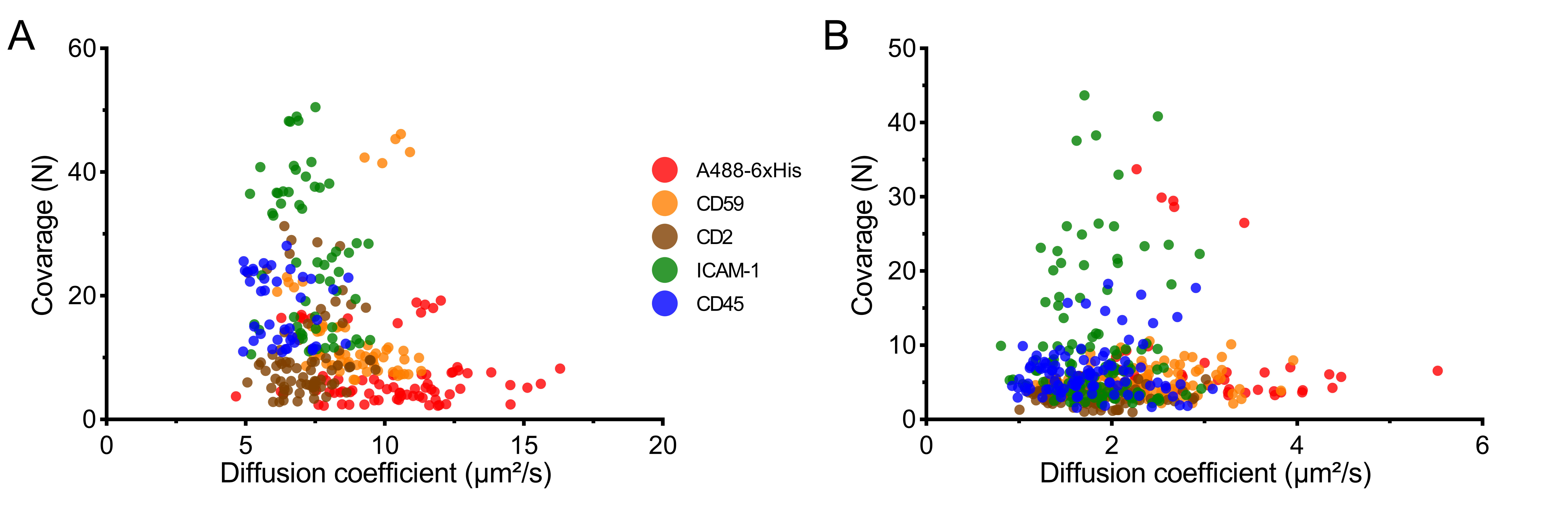

**Supp Fig. 7. Number of molecules (coverage) vs diffusion time.** Membrane coverage correlated to the diffusion coefficients obtained during the experiments for (A) POPC and (B) DPPC:Chol in GUVs. The plot shows no correlation between the mobility and membrane coverage, displaying that the experiments were performed under non-crowding affected conditions.

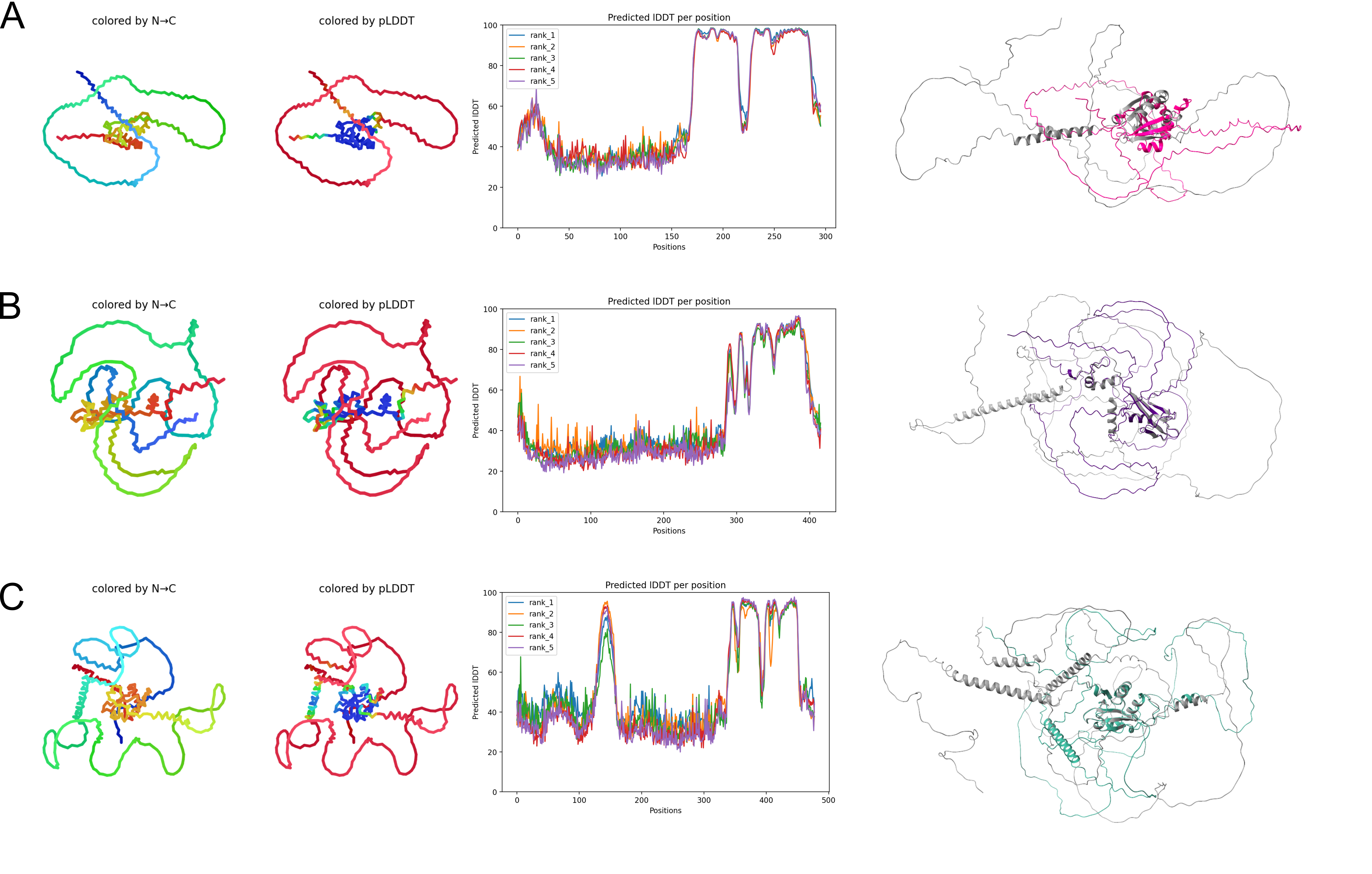

**Supp. Fig. 8. ECD Structural models via ALPHAFOLD2.** Model obtained via Collabfold, multiple sequence alignment performed via Jackhmmer, displayed is the highest ranked out of 5 models performed, with the LDDT graph for all models displayed next to the structures. Also shown are alignments to published structures for the whole protein’s published Alphafold prediction. (A) CD34, AlphaFold: AF-P28906-F1, (A) CD34, AlphaFold: AF-P28906-F1, (B) PODXL, AlphaFold: AF-O00592-F1, (C) PODXL2, AlphaFold: AF-Q9NZ53-F1. Some of the structures cannot be resolved with Alphafold, presumably due to glycosylation.

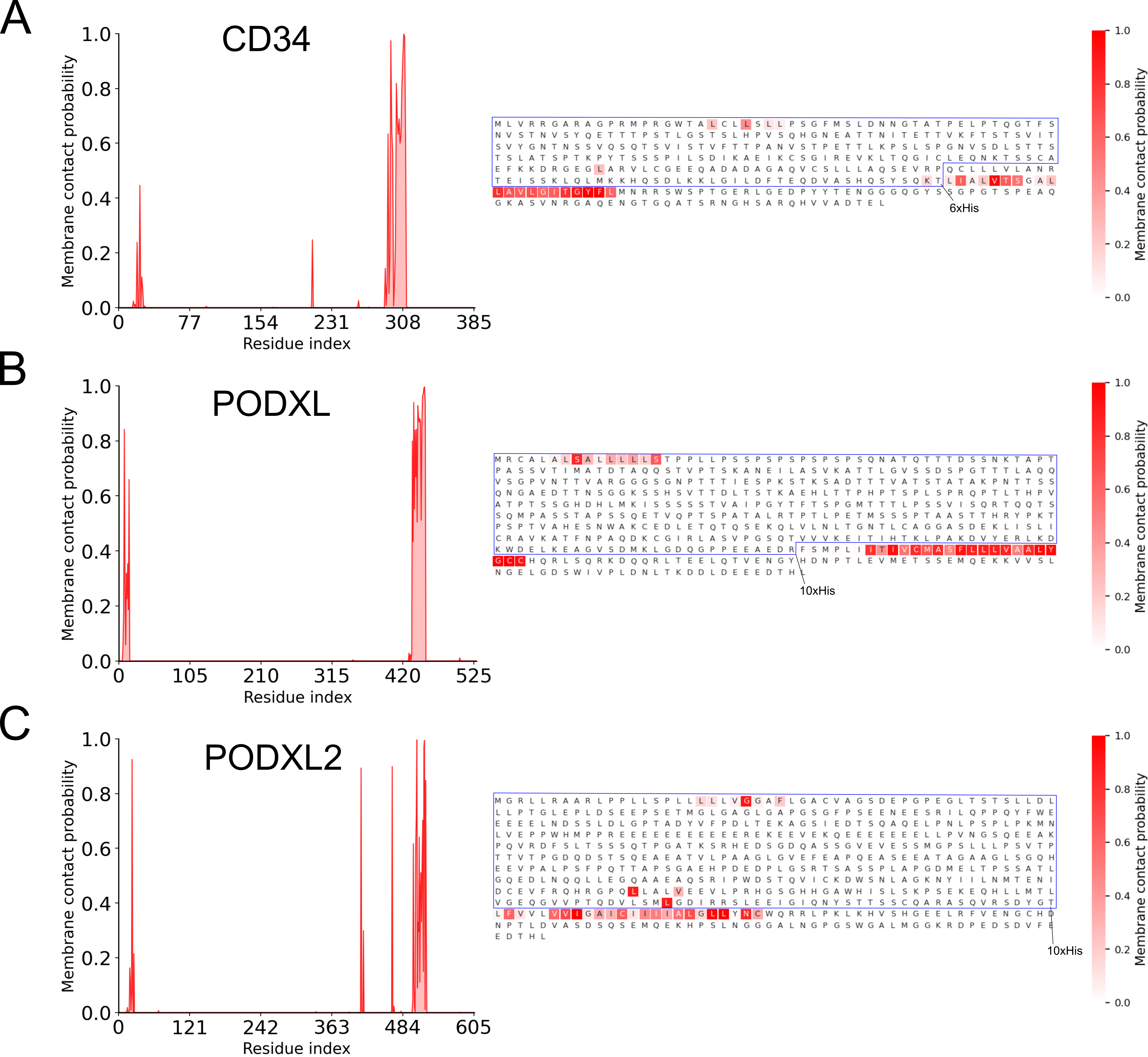

**Supp. Fig. 9. Membrane contact probability.** The analysis was done on the full-length proteins, and blue frames show extracellular domain of the proteins used in the study. The 6xHis-tag or 10xHis-tag at C-terminus is shown with arrow for each protein.

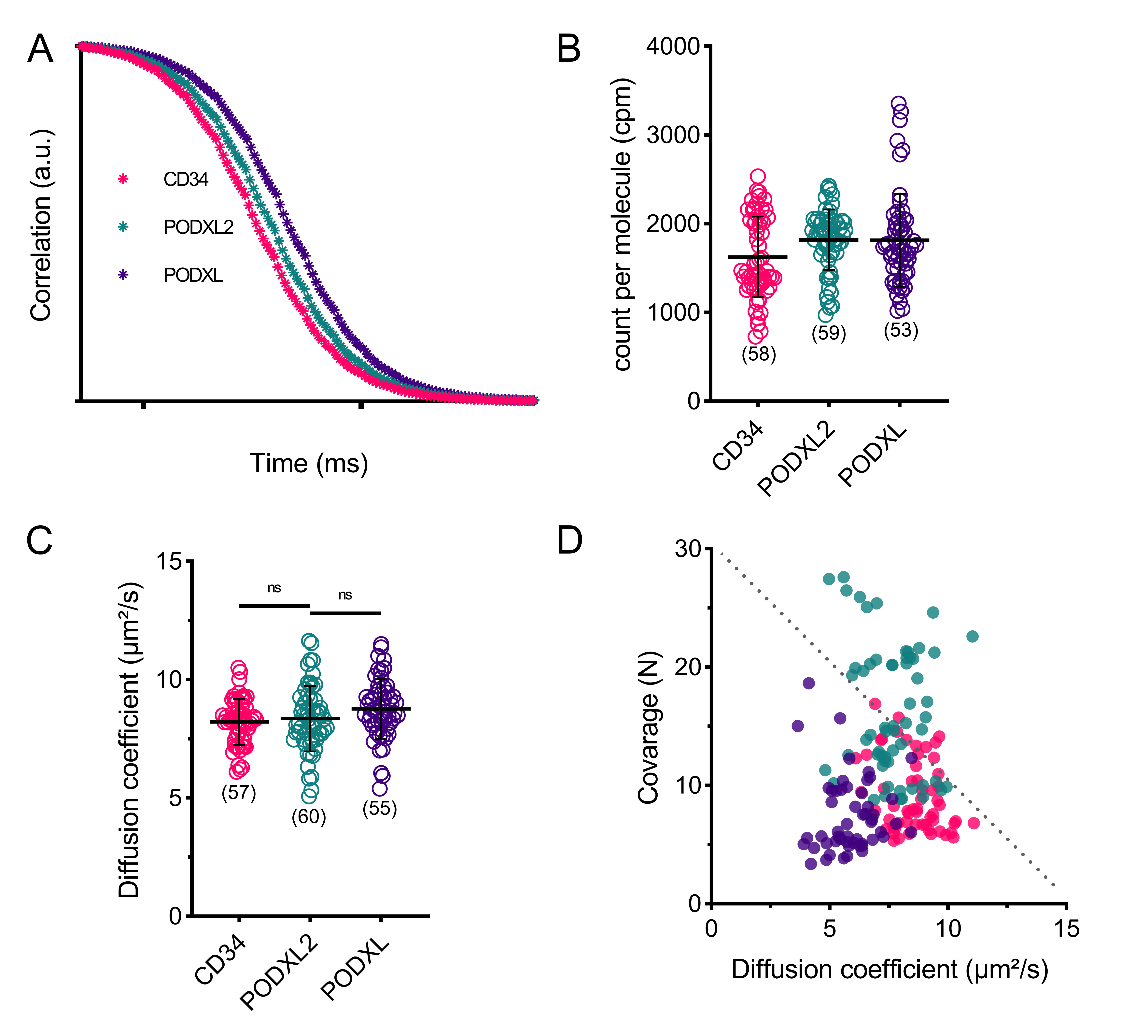

**Supp. Fig. 10. Control experiments for CD34, PODXL2, and PODXL.** (A) FCS curves. (B) Count per molecule for the FCS measurement in GUVs. (C) Diffusion of the membrane dye (AbStR-DPPE) in GUVs. (D) Number of molecules (coverage) vs diffusion time. Non-parameterized ANOVA was used to determine significance (ns: non-significant). Horizontal lines show the mean and error bars represent the standard deviation. The number of data points obtained from experiments are indicated on the graphs in parentheses.

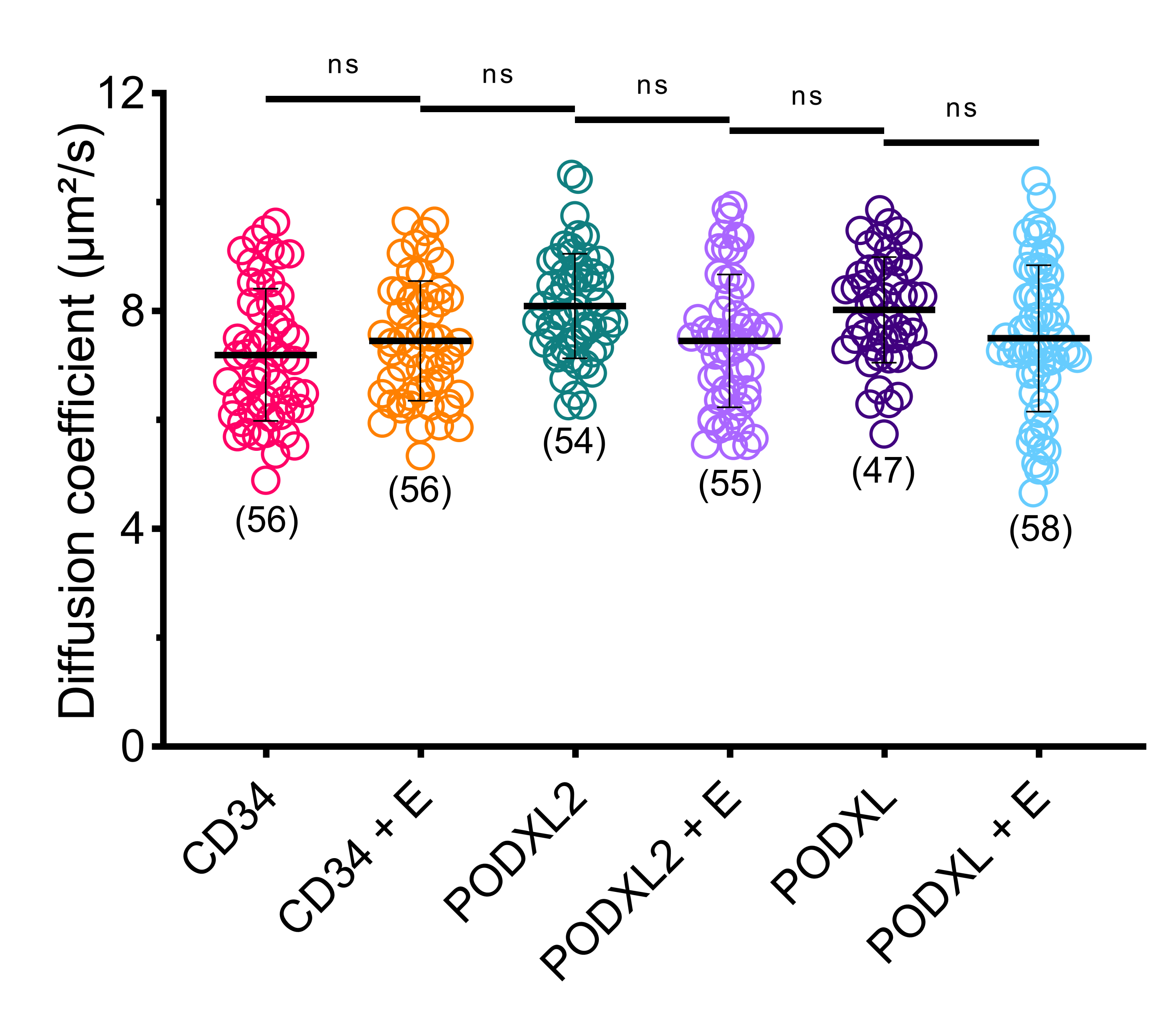

**Supp. Fig. 11. Diffusion of membrane dye (AbStR-DPPE) in GUVs.** Non-parameterized ANOVA was used to determine significance (ns: non-significant). Horizontal lines show the mean and error bars represent the standard deviation. The number of data points obtained from experiments are indicated on the graphs in parentheses. E stands for the enzyme treatment for the deglycosylation.

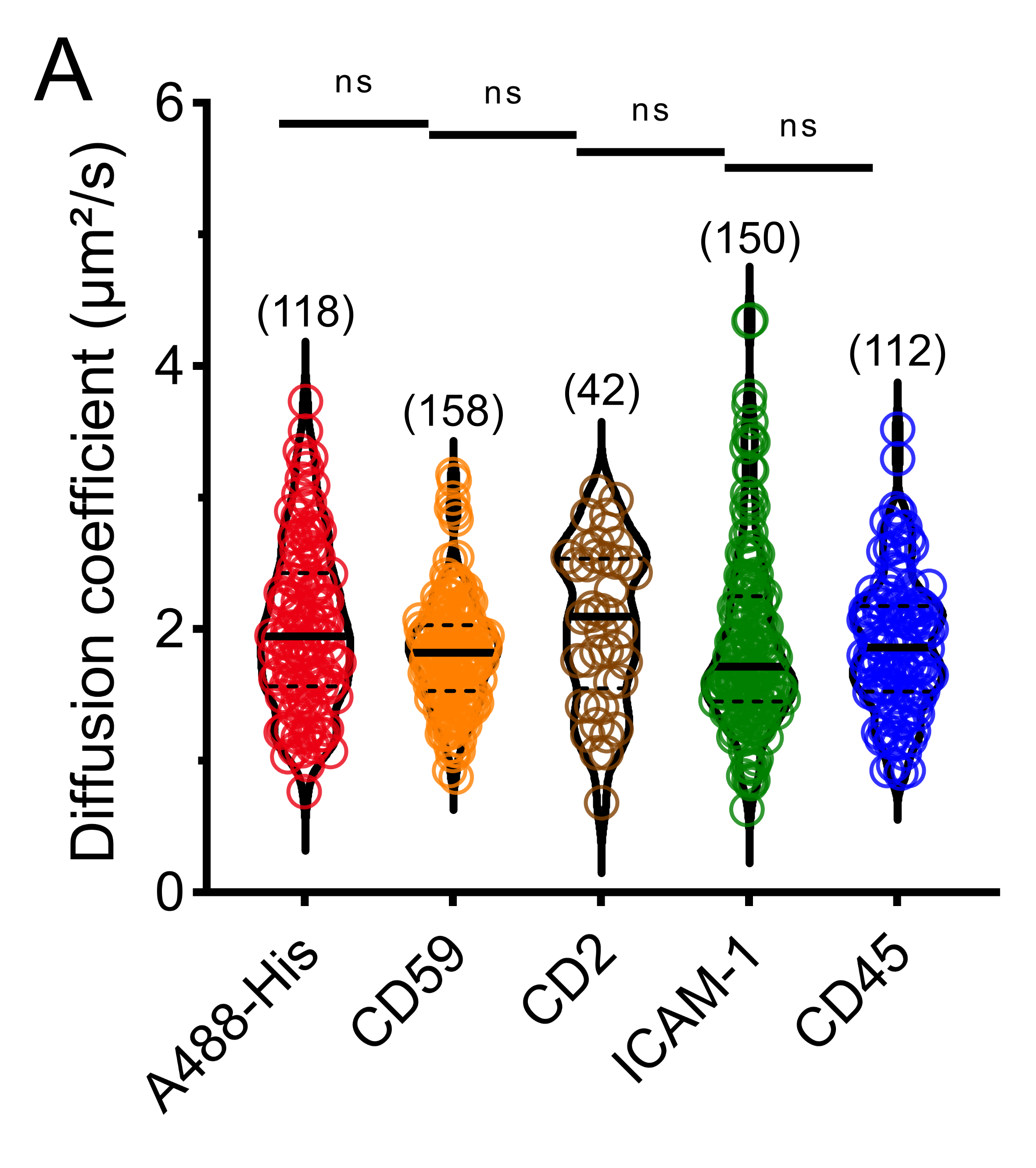

**Supp. Fig. 12. Diffusion of membrane dye (AbStR-DPPE) in GPMVs.** Non-parameterized ANOVA was used to determine significance (ns: non-significant). Horizontal thick lines show the median and dotted lines represent the quartiles. The number of data points obtained from experiments are indicated on the graphs in parentheses.

**Supplementary Tables**

**Supp. Table 1: Molecular weight and required values for dye volume calculation for each of the ECDs investigated.**

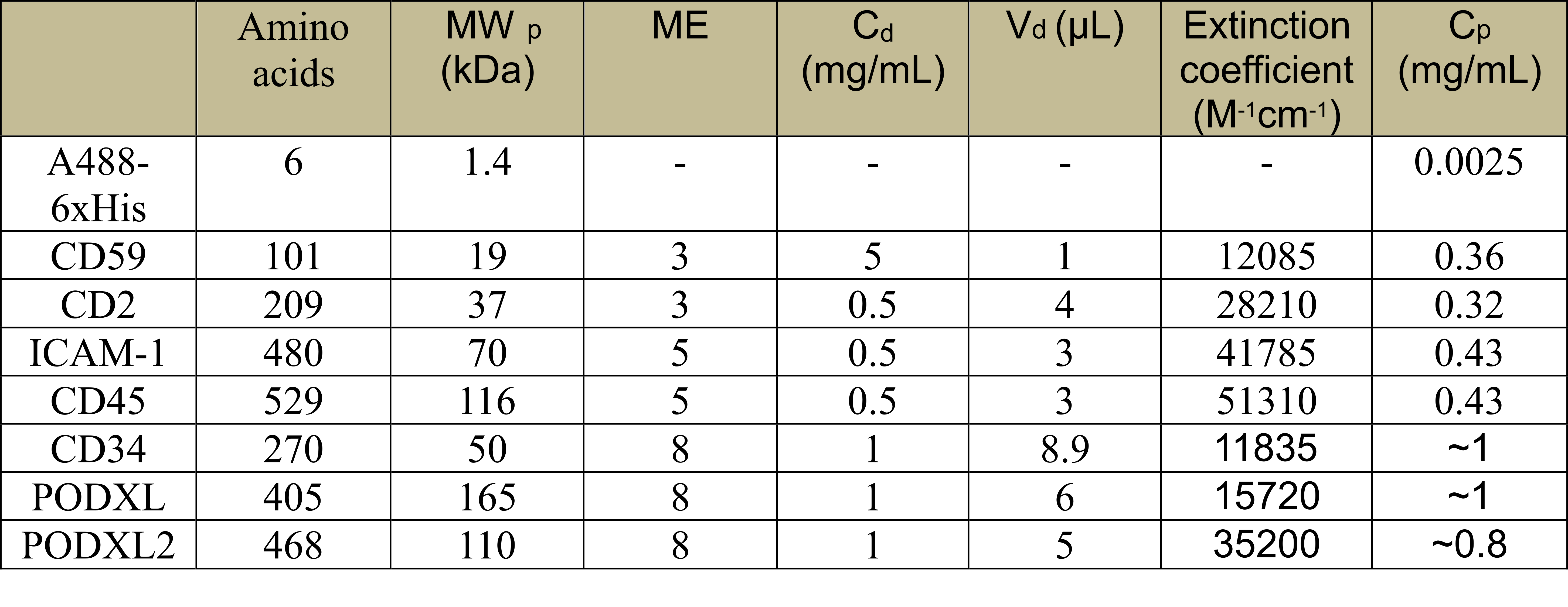

**Supp. Table 2: Alphafold2 (colabfold) parameters chosen during the model generation.**

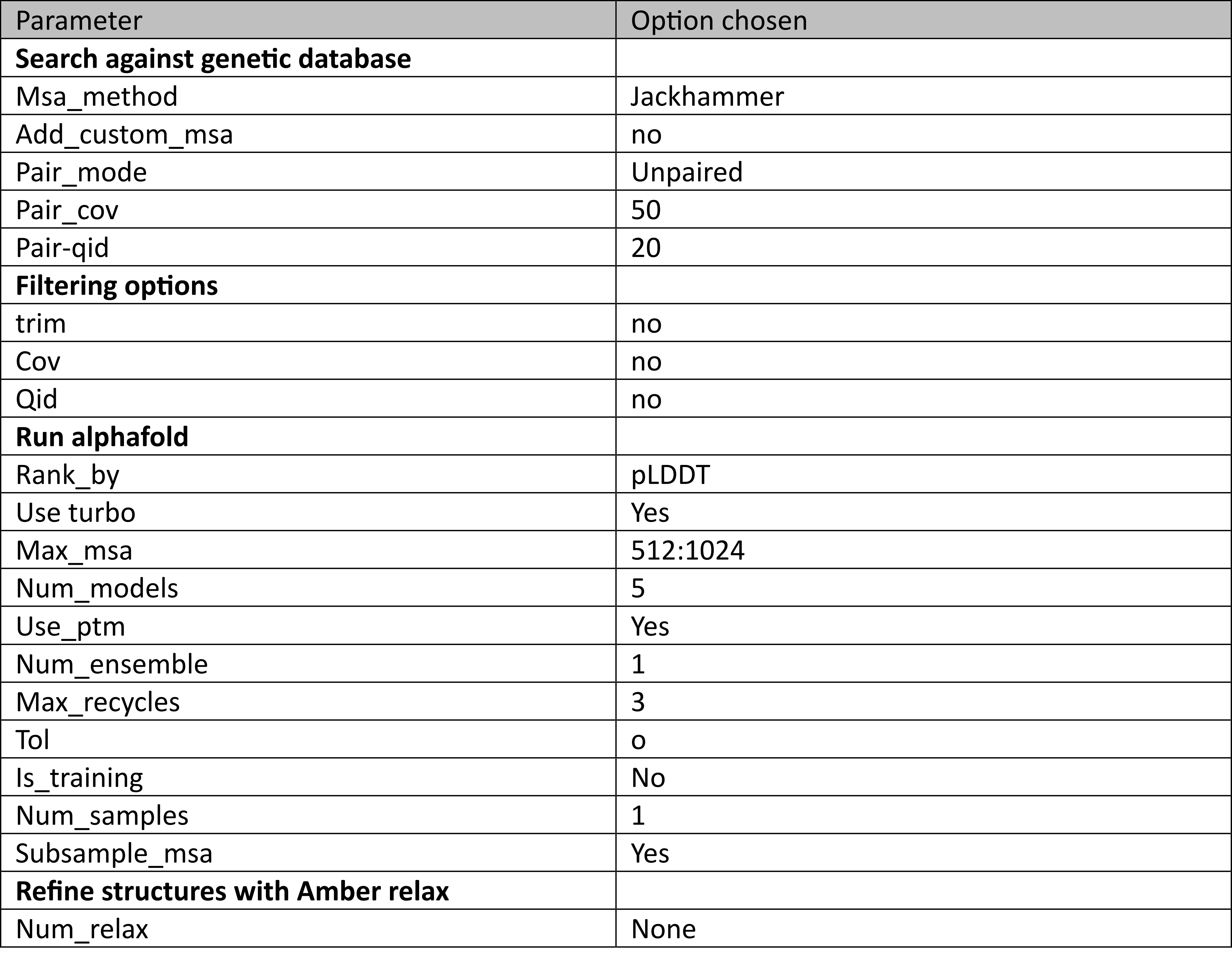

**Supp. Table 3: Amino acid sequences of the ECDs used in the project.** Grey colors show the N-terminal 6xHis-Tag or 10xHis-tag.

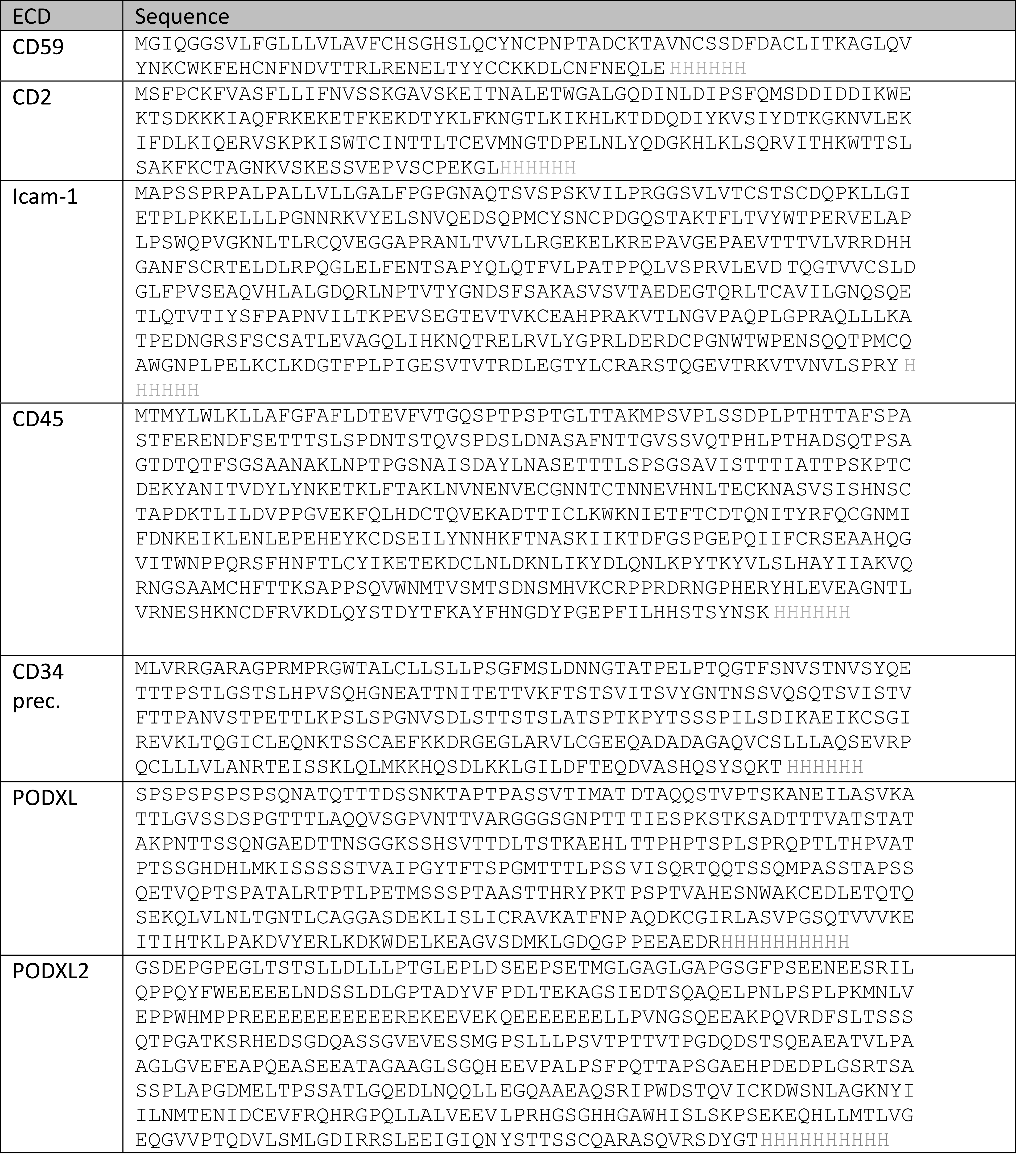
